# Admixture and environment shape population genetic and phytochemical variation across a conifer hybrid zone

**DOI:** 10.1101/2025.07.31.667945

**Authors:** Kathryn A. Uckele, Casey S. Philbin, Lora A. Richards, Lee A. Dyer, Joshua P. Jahner, Thomas L. Parchman

## Abstract

Ancestry variation in hybrid zones can reflect the causes and genetic basis of reproductive isolation and result in novel phenotypic variation with the potential for extended ecological effects. Junipers (*Juniperus*) are foundational tree species in many semi-arid landscapes of western North America and often hybridize in zones of secondary contact. Such hybridization can be ecologically significant in foundational tree species, due to the strong genetic control and ecological consequences of plant chemistry. We generated genetic and phytochemical data to analyze hybridization among *Juniperus grandis*, *J. occidentalis*, and *J. osteosperma* in western Nevada and its impact on plant chemistry. We used population genomic data (9,125 SNPs; 326 individuals; 25 populations) to quantify patterns of genetic variation across populations and species and characterize ancestry variation in hybrids. While populations within species showed little genetic differentiation, the parental species formed distinct, monophyletic lineages with clear phenotypic and ecological differences. Hybrids occupied intermediate environments, contained ancestry from all three parents, and were mainly F_1_ or backcross hybrids. Phytochemical data (GC-MS; 163 terpenoid compounds) were likewise analyzed to understand the consequences of hybridization for plant chemistry. The parental species and hybrids displayed distinct phytochemical profiles, with hybrids often characterized by a combination of transgressive and intermediate chemical concentrations. Our results illustrate that geography and environment shape hybrid ancestry for a syngameon involving three *Juniperus* species, and that admixture generates novel phytochemical variation likely to have ecological consequences.

## 2. Introduction

Genome-scale sequencing has increasingly uncovered signatures of hybridization across diverse taxa, revealing key roles in diversification (Soltis et al. 2015; Tank et al. 2015; Alix et al. 2017), domestication (Arnold 2004), and the evolution of invasive species (Ellstrand and Schierenbeck 2000). The proximate and ultimate outcomes of secondary contact depend on stochasticity, ecological context, and reproductive barriers (Baack and Rieseberg 2007; Mandeville et al. 2017; McFarlane et al. 2024). If reproductive isolation is nearly complete, F_1_ hybrids may have reduced fitness, causing F_2_ and backcross hybrids to be rare (Burke and Arnold 2001; Burton et al. 2013). Alternatively, without strong barriers, recombinant hybrids may form, allowing introgression among parental species, and offering insights into the genetic basis of speciation and complex trait genetics (Buerkle and Lexer 2008; Gompert et al. 2017). Hybridization may also contribute to adaptation and diversification by leading to adaptive introgression and generating novel phenotypic variation in hybrids (Rieseberg et al. 1999, 2003, 2007; Kagawa and Takimoto 2018).

Admixture among plant species can dramatically shape phenotypic variation, with important ecological and evolutionary consequences. Biologically active mixtures of plant specialized metabolites, a key phenotypic axis, mediate species interactions (Wimp et al. 2005), structure ecological communities (Abrahamson et al. 2003; Poelman et al. 2009), and regulate the flow of resources through ecosystems (Schweitzer et al. 2008). Plant hybridization often generates novel phytochemical variation with the potential to shape interactions and community dynamics. In many cases, hybridization produces transgressive variation, where chemical traits exceed parental ranges, expanding the variation available for selection (Rieseberg et al. 1999, 2003, 2007). *Juniperus* is a genus characterized by high diversity of specialized metabolites in both leaves and cones, including flavonoids, coumarins, lignans, tropolones, and the production of essential oils rich in terpenoids such as α-pinene, sabinene, and cedrol (Schwartz et al. 1980; Keeling and Bohlmann 2006; Seybold et al. 2006; Markó et al. 2011; Pardikes et al. 2019). These compounds likely play significant roles in ecological interactions, but most phytochemical investigations for this genus have focused on applied uses, such as antibacterial properties, effects on livestock grazing, and medicinal potential (Bozyel et al., 2024). Even though Great Basin *Juniperus* species house a rich community of arthropods (Pardikes et al. 2019) that are likely important to vertebrate communities supported by juniper woodlands, there is much to be learned about how the rich chemistry of these trees mediates interactions with arthropods and vertebrates. Simple questions about how metabolomic profiles vary predictably across species and also mirror genetic structure are an obvious starting point for characterizing these chemically mediated interactions in a dominant ecosystem across the US Western landscape. Here we quantified genetic and chemical variation across a juniper hybrid zone as a basis for further understanding of how admixture may generate novel traits, influence species interactions, and shape broader ecological and evolutionary processes (Volf et al. 2024).

*Juniperus* trees and shrubs are foundational woody plants in many arid regions of the Northern Hemisphere. The serrate leaf junipers of North America (21 species) diversified into the arid and mountainous regions of the western U.S. and Mexico over the past ∼23 million years (Mao et al. 2010; Uckele et al. 2021). Hybridization is common in this clade (Adams 1994; Adams et al. 2017, 2020) and has been documented among the focal serrate junipers of this study: *J. grandis*, *J. occidentalis*, and *J. osteosperma* (Terry et al. 2000; Terry 2010). *Juniperus grandis* (previously *J. occidentalis* var. *australis*) and *J. occidentalis* (previously *J. occidentalis* var. *occidentalis*) are sister species that diverged ∼5 mya (Uckele et al. 2021). *Juniperus grandis* grows on dry rocky slopes within the eastern Sierra Nevada of California, and *J. occidentalis* occurs on dry rocky foothills and mountain slopes in northern California, eastern Oregon, and southern Washington (Fig. 1). Basal to these two species is *J. osteosperma*, whose lineage diverged ∼11 mya and currently occurs across much of the Great Basin and Colorado Plateau (Fig. 1; Adams 2014). These species can form large stands in otherwise treeless regions, often coexisting with pinyon pines (*Pinus* subsection *Cembroides*) to create key ecosystems in the western U.S. and Mexico (Miller and Wigand 1994; Weisberg et al. 2007). Pinyon-juniper woodlands, found in drier climates than many other tree-dominated ecosystems, support a high biodiversity of plants, birds, mammals, and invertebrates (Gottfried et al. 1995; Tonkel et al. 2021).

**Figure 1.**
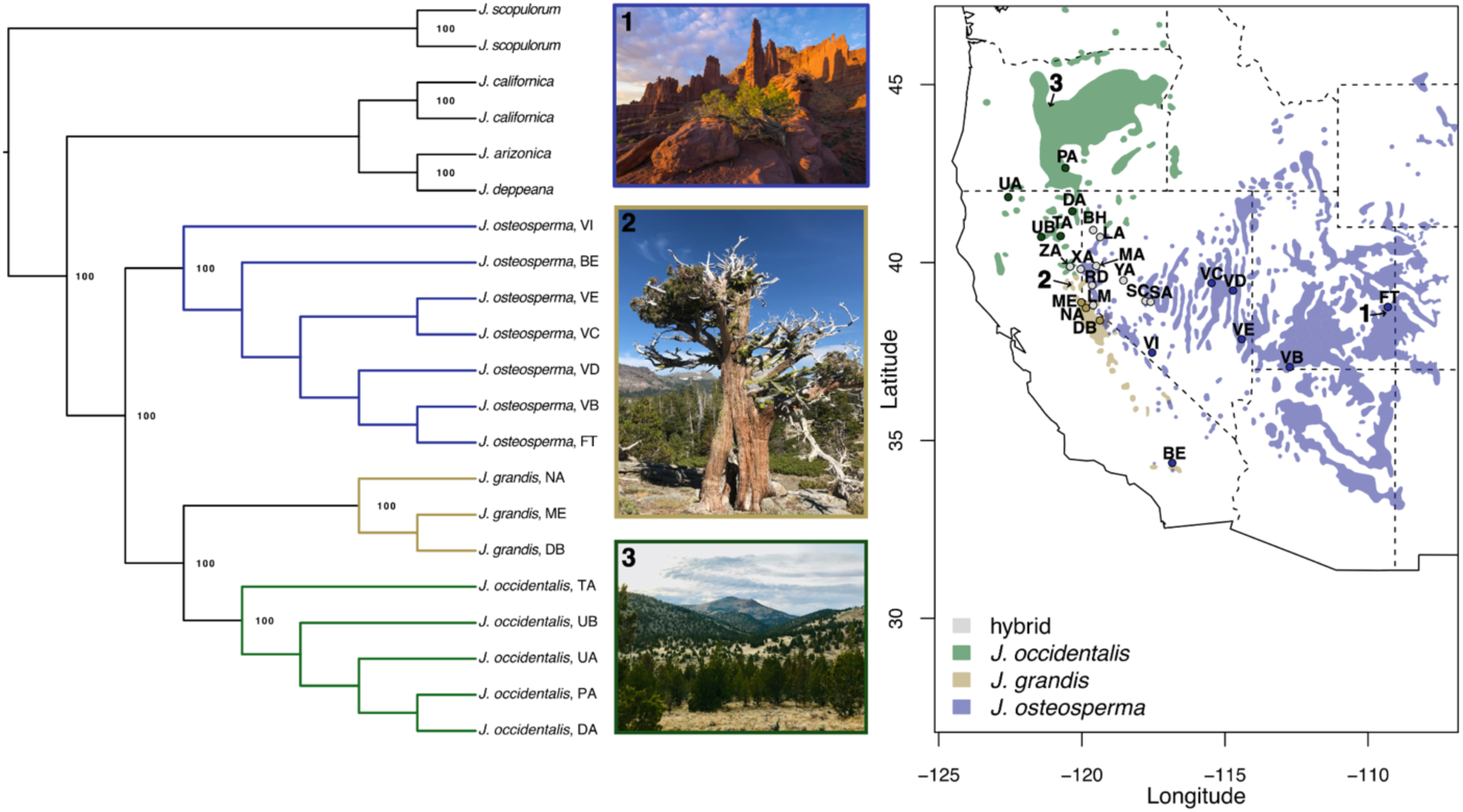
The phylogenetic tree on the left shows evolutionary relationships among the parental taxa, with outgroup taxa indicated by black tips. Branches and tips of each parental lineage are colored according to the sampling localities on the map to the right. Representative photos of the parental taxa appear between the tree and the map: (1) prostrate *Juniperus osteosperma* at Fisher Towers, UT (photo credit: Floris van Bruegel), (2) *J. grandis* at Donner Lake, CA, and (3) a *J. occidentalis* forest near Smith Rock State Park, OR. The map on the right displays each sampling locality (labeled with its two-letter code), overlaid on colored polygons representing the ranges of the parental species (*J. occidentalis* in green, *J. grandis* in gold, *J. osteosperma* in blue) and hybrid populations in gray. Approximate photo locations are also labeled on the map. Additional locality information is provided in Supplementary Table 1.

During the Pleistocene glaciation cycles, the ranges of many North American tree species, including junipers, shifted in response to climate changes (Roberts and Hamann 2015). At the height of the Wisconsin glacial (ca. 25–21 Kya), *J. grandis*, *J. occidentalis*, and *J. osteosperma* likely occupied isolated glacial refugia. Analyses of packrat midden macrofossils suggest that *J. grandis* occurred in the mesic western aspect of the central Sierra Nevada range, *J. osteosperma* expanded into areas significantly south of its contemporary range in the current-day Sonoran, Mojave, and Chihuahuan deserts, and *J. occidentalis* occurred, at least temporarily, in the northern Great Basin (Betancourt et al. 1990). As the last glacial period ended (∼ 12 Kya), *J. osteosperma* began migrating north, eventually entering into secondary contact with *J. grandis* and *J. occidentalis* in the Sierra Nevada foothills. The resulting hybrid zone in western Nevada spans the Sierra Nevada-Great Basin ecotone (Vasek 1966), a transitional region where hybrid zones frequently cluster (Remington 1968; Swenson and Howard 2005). Previous studies using morphology, phytochemistry, and genetic markers identified this hybrid zone and detected introgression into the *J. osteosperma* background but lacked the genetic resolution to analyze hybrid zone composition and fine-scale genetic variation (Vasek 1966; Terry et al. 2000, 2010; Adams 2013a,b). Genomic data stands to clarify the factors shaping reproductive isolation and admixture and provide insights into the phenotypic consequences of ancestry variation in hybrids.

We leveraged genetic, phytochemical, and environmental variation across three juniper species and their hybrids to address four key objectives: 1) resolve the evolutionary history of the parental species, 2) characterize ancestry variation in hybrids, 3) examine the relationship between ancestry and environmental variation, and 4) assess hybridization’s impact on phytochemistry. Using genomic and phytochemical data from a large sample of individuals and populations, we found that hybrids, primarily F_1_s and backcrosses, possess ancestry from all three parental species, with hybrid ancestry varying along geographic and environmental gradients. The hybrid zone spans an altitudinal gradient associated with intermediate climatic conditions, suggesting a role for local adaptation in shaping ancestry patterns. Additionally, hybrids exhibit distinct phytochemical profiles, with the potential to shape novel community structures and interactions. Our findings contribute to an understanding of the factors that shape admixture and its consequences for phytochemical variation in a foundational tree syngameon and provide a foundation for exploring the ecological consequences of this variation.

## 3. Methods

### 3.1 Sampling, library preparation, and DNA sequencing

We collected leaf samples from 386 individuals for genetic and phytochemical analyses. Samples were distributed across three *J. grandis* populations (N = 37), five *J. occidentalis* populations (N = 56), seven *J. osteosperma* populations (N = 91), and ten hybrid populations (N = 142). An additional 60 samples from outgroup taxa (*J. arizonica*, *J. californica*, *J. deppeana*, and *J. scopulorum*) were used for phylogenetic analyses (Supplementary Table 1).

DNA was extracted with Qiagen DNeasy Plant Mini Kits (Qiagen Inc., Valencia, CA, USA) and quantified with a QIAxpert microfluidic analyzer (Qiagen Inc., Valencia, CA, USA). Sequencing libraries were prepared following the ddRADseq protocol described by Parchman et al. (2012). Briefly, DNA was digested with *Eco*RI and *Mse*I restriction enzymes, custom oligos with Illumina adapters were ligated to fragment ends, and fragments were PCR amplified using a high-fidelity proofreading polymerase (Iproof polymerase, BioRad Inc., Hercules, CA, USA) before being pooled for sequencing. Libraries were size-selected (350–450 bp) using the Pippin Prep System (Sage Sciences, Beverly, MA, USA) at the University of Texas Genome Sequencing and Analysis Facility (Austin, TX, USA). Four lanes of single-end 100 bp sequencing were performed on an Illumina HiSeq 2500 platform at the University of Wisconsin-Madison Biotechnology Center (Madison, WI, USA). Contaminant reads (Illumina adapters, primers, PhiX, and *E. coli*) were removed using a filtering pipeline (https://github.com/ncgr/tapioca), and demultiplexing was performed with custom scripts. The sequencing data are available on the NCBI Sequence Read Archive under BioProject ID PRJNA1037072.

### 3.2 Phylogenetic analyses

To clarify the evolutionary relationships among the parental species, phylogenetic inference was performed on pure parental and outgroup samples (one sample per population). Reads were *de novo* assembled and filtered with `iPyRAD` (v.0.7.19; Eaton and Overcast, 2020), with detailed methods provided in the Supplementary Materials. Phylogenetic analysis was conducted with `TETRAD` (Eaton et al. 2017), which implements the SVDquartets algorithm (Chifman and Kubatko, 2014; 2015). All possible species quartets were analyzed to infer the unrooted species tree, which was manually rooted with the *J. scopulorum* outgroup. Node support values were estimated based on 100 bootstrap replicates.

### 3.3 Genetic variation across populations, species, and hybrids

A *de novo* reference assembly of 160,256 contigs was generated by clustering unique reads present in at least four samples with `cdhit` (v. 4.6; Li and Godzik, 2006; Fu et al. 2012) according to a similarity threshold of 80%. Reads were then aligned to this reference using the `aln` and `samse` algorithms in `bwa` (v. 0.7.5a; Li and Durbin 2009). Variants were called and filtered using `samtools` (v. 1.3; Li and Durbin 2009), `bcftools (`v. 1.3; Li 2011; Danecek et al. 2021), and `vcftools` (v. 0.1.14; Danecek et al. 2011). Biallelic SNPs were filtered using several quality criteria: a site quality (QUAL) threshold >20, genotype quality (GQ) >8, and a minor allele frequency >0.05. Additionally, SNPs were retained only if they were present in at least 60% of samples (i.e., maximum 40% missing data), and the dataset was thinned to include one SNP per locus. Potentially paralogous loci were removed using the `HDplot` pipeline for genotyping-by-sequencing data (McKinney et al. 2017). Separate variant calling procedures were conducted for 1) all 386 samples (including outgroups) and 2) a subset of 326 samples comprising the target parental species and hybrids.

Individual ancestry proportions were inferred with `entropy` (Gompert et al. 2014; Shastry et al. 2021), a Bayesian clustering method analogous to `STRUCTURE` (Pritchard et al. 2000; Falush et al. 2003). In addition to estimating ancestry proportions (*q*), `entropy` leverages genotype likelihoods and population allele frequencies to infer genotype probabilities while accounting for uncertainty, which were used in all subsequent analyses requiring genotype information. For the analysis of all samples including outgroups, we set the number of source populations (*K*) to seven, corresponding to the number of species sampled. For the analysis of parental species and hybrids, *K* was set to three, reflecting the number of parental species. To facilitate convergence, we initialized the MCMC chains with cluster membership probabilities derived from linear discriminant analysis and *K*-means clustering of the first five principal components calculated from genotype likelihoods (see Jombart et al. 2010). Four independent chains were run with a burn-in phase of 30,000 iterations, followed by sampling every tenth iteration for 60,000 iterations. Convergence and mixing were assessed visually with R (v. 4.1.2; R Core Team 2020).

`Entropy` was also used to estimate interspecific ancestry, defined as the proportion of an individual’s genome where alleles were inherited from different source populations (Shastry et al. 2021). By comparing individual ancestry proportions (*q*) to interspecific ancestry proportions (*Q*), it is possible to distinguish ancestry classes, such as F_1_, F_2_, backcrosses, and later-generation hybrids. For example, both F_1_, F_2_, and later-generation hybrids are expected to have individual ancestry proportions of 0.5. However, F_1_ individuals have *Q* values of one because each copy of their genome is inherited from a different parental species, whereas later-generation hybrids have *Q* values less than one due to recombination breaking up long tracts of high interspecific ancestry (Gompert et al. 2014; Shastry et al. 2021). For *K* values greater than two, the interpretation of ancestry classes becomes more complex due to the increased number of possible interspecific ancestry combinations. To address this, we treated *J. grandis* and *J. occidentalis* as a single ancestral population, given their much lower divergence from one another compared to their combined divergence from *J. osteosperma* (see Results).

To visualize genetic structure across populations and species, principal components analysis (PCA) was performed on the genotype probabilities. To mitigate the effects of unbalanced sampling of species, we randomly selected 15 individuals from each species for the analysis of all samples. To quantify genetic differentiation among populations, Nei’s *D* (Nei 1972) and Hudson’s F_ST_ (Hudson et al. 1992) were calculated from population allele frequencies with custom scripts in `R`. A neighbor-joining tree was constructed from Nei’s *D* values using the *nj* function (`ape` package; Paradis and Schliep 2018). Mantel tests (`vegan` package; Oksanen et al., 2020) using Spearman correlations and complete permutation enumeration assessed the relationship between genetic (Nei’s *D*) and geographic distances for *J. osteosperma* and *J. occidentalis* (*J. grandis* was excluded due to insufficient population sampling). Geographic distances between populations were calculated from geographic coordinates (Supplementary Table 1) using the *earth.dist* function (`fossil` package; Vavrek 2011).

We calculated two estimates of genome-wide genetic diversity – Θπ (the average number of polymorphisms per pairwise sequence comparison; Tajima 1983) and Θw (the number of segregating sites; Watterson 1975) – for each population using methods that account for genotype uncertainty, as implemented in `ANGSD` (v. 0.923; Korneliussen et al. 2014). First, we estimated the site allele frequency likelihood (SAF) from the BAM files and *de novo* reference assembly using the *-doSaf* option. We then used the `realSFS` utility (Nielsen et al. 2012) to estimate the folded site frequency spectrum (SFS) from the SAF and calculated Θπ and Θw for each site using the *saf2theta* option. To evaluate whether the populations are evolving neutrally and conforming to mutation-drift balance, we calculated Tajima’s *D* (Tajima 1989) for each site using the `thetaStat` utility and the *doStat* option (Korneliussen et al. 2013).

### 3.4 Ecological divergence among parental taxa and hybrids

To assess divergence in the contemporary climate envelopes of each parental species range and the hybrid zone, we analyzed range-wide climate data with discriminant analysis of principal components (DAPC; Jombart et al. 2010). For each parental species, 200 geographic coordinates were randomly selected from plot observations in the US Forest Service Forest Inventory and Analysis (USFIA) database (http://www.fia.fs.fed.us/). Since hybrids were not specifically identified in the database, we classified all *J. osteosperma* plot observations west of 118° W longitude as hybrids, based on the geographic distribution of hybrid ancestry (Fig. 4b). For each geographic coordinate, 25 climate variables were extracted at 30-arcsec (approximately 800 meters) resolution from PRISM (PRISM Climate Group, 2007) and the Climate Water Deficit Toolbox (Dilts and Yang, 2015). Using species and hybrid assignments as *a priori* groups, we retained 10 principal components of scaled climate data and performed linear discriminant analysis with the *dapc* function (*adegenet* package; Jombart, 2008) in `R`.

To assess the extent to which climate predicts ancestry variation across our sampled populations, we used random forest models. For each parental and hybrid population, we extracted 30 climate variables from their geographic coordinates using the same method as the range-wide analysis described above. The model assigned each individual to an ancestry group (*J. grandis*, *J. occidentalis*, *J. osteosperma*, or hybrid) based on the climate data associated with their population of origin. The classification model, implemented with the `randomForest` package in `R` (Liaw and Wiener 2002), was trained on 70% of the individuals and tested by predicting group membership for the remaining 30%. We developed a second random forest model using the `H2O` framework in `R` (LeDell et al., 2020) to predict hybrid ancestry proportions (*q*) from 30 climate and two geographic predictors (latitude and longitude). The dataset was randomly split into 80% training and 20% test sets, and hyperparameters were optimized using a random grid search with leave-one-out cross-validation. The best model, selected based on the lowest cross-validated mean squared error, was evaluated using root mean squared error (RMSE), mean absolute error (MAE), and R^2^ on both cross-validation and test sets.

To assess the relationship between genetic variation and climate variation, we performed redundancy analysis (RDA; Legendre and Makarenkov, 2002; Capblancq and Forester, 2021) using genotype probabilities as the response matrix. To minimize multicollinearity among predictor variables, we sequentially removed climate variables with the highest mean correlations until all remaining variables had correlations below 0.7. The resulting set of seven climate variables was then scaled and used as multivariate predictors. RDA was conducted with the *rda* function (`vegan` package) in `R`.

### 3.5 Phytochemical variation across species and hybrids

We quantified specialized metabolite profiles for 314 of 326 focal individuals. Silica-dried leaf samples were lysed using a Qiagen TissueLyser II (Qiagen Inc., Valencia, CA, USA), and 20 mg of tissue was extracted with 1 mL hexanes containing an internal standard of n-Eicosane (0.25 mM). After vortexing, samples were sonicated in ice water for 15 minutes, centrifuged, and chilled at -20°C to minimize evaporation before transfer to GC-MS autosampler vials. Extracts (1 μL) were injected onto an Agilent 7890A gas chromatograph coupled to an Agilent 5975C quadrupole mass spectrometer with an Agilent HP-5MS capillary column (Agilent Technologies Inc., Santa Clara, CA, USA). Full GC-MS protocol and data processing details are in the Supplementary Materials. The final phytochemical dataset included concentrations for 163 terpenoid compounds, 56 of which (34%) were annotated using a NIST match (score > 60) against the Adams GCMS library (Adams, 2007). When a compound matched multiple peaks, the peak with the closest retention index was annotated as that compound, and the other peaks were categorized as structural analogs.

PCA was performed to examine variation in terpenoid compound profiles. Due to strong positive skew in the GC-MS data, a log2 transformation was applied before analysis. Preliminary PCA identified 14 outliers from the ZA population (*J. grandis x J. occidentalis* hybrids) that dominated variance in PCs 1 and 2; these were excluded in subsequent analyses, leaving 300 samples. PCA was conducted using the *prcomp* function in `R` with centering but no scaling. To assess whether terpenoid variation could classify individuals into ancestry groups, we performed a random forest analysis using terpenoid data with the `randomForest` package (Liaw and Wiener, 2002) in `R`. The model was trained on 70% of the data and evaluated on the remaining 30%.

To categorize terpenoid compounds as dominant, intermediate, or transgressive, we used pairwise Wilcoxon rank-sum tests with the Holm-Bonferroni correction (*pairwise.wilcox.test* in `R`). Due to positive skew, we present the non-parametric test results, though one-way ANOVA with Tukey’s HSD produced similar results. Compounds were classified as dominant if the hybrid group was significantly different from at least one parental species but not all. Intermediate compounds were those where the hybrid group differed from both parental species but had concentrations within the parental range. Transgressive compounds were those where the hybrid group differed significantly from all parental species, with mean concentrations outside the parental range. We also quantified the number of transgressive hybrids exceeding parental ranges and calculated total terpenoid concentrations by summing all compounds, assessing group differences with pairwise Wilcoxon tests.

To disentangle genetic, geographic, and environmental influences on phytochemical variation, we performed variance partitioning using partial redundancy analysis (pRDA). The response matrix included terpenoid concentrations and the predictor matrix included: 1) the first two principal components from PCA of climate data (explaining 73.74% of variance), 2) the first two principal components from PCA of genotype probabilities (50.56% of variance), and 3) geographic coordinates (latitude and longitude). A standard RDA was first conducted to estimate total constrained variance. Three pRDAs were then performed, conditioning on two predictor sets at a time, to isolate the unique contributions of climate, genetics, and geography. Results were consistent with variance partitioning using the *varpart* function (`vegan` package) in `R`.

To examine terpenoid diversity and variation within and among ancestry groups, we used diversity indices and distance-based analyses. To account for uneven sample sizes, we rarefied the data by randomly selecting six samples per population for *J. osteosperma* (for a total of 36 samples), seven per population for *J. occidentalis* (for a total of 35 samples), and four per hybrid population (for a total of 40 samples). All 36 available samples for *J. grandis* were included. Alpha diversity (per sample) and gamma diversity (per ancestry group), expressed as the effective number of species, were calculated by exponentiating the Shannon diversity index (`vegan` package) separately for terpenoid concentration data and presence/absence data.

A generalized linear model with a Gaussian distribution was used to test differences in mean alpha diversity among ancestry groups, followed by post-hoc Tukey’s HSD tests (`multcomp` package; Hothorn et al., 2008). Additive beta diversity was also calculated across individuals for each ancestry group, with detailed methods provided in the Supplementary Materials.

We employed two distance-based analyses to evaluate beta diversity within and between ancestry groups: PERMANOVA to test for differences in terpenoid composition and a multivariate analogue of Levene’s test to assess homogeneity of variances. Since PERMANOVA is sensitive to variance differences, testing for multivariate homogeneity of variances ensured consistent variances across groups while providing additional insight into beta diversity. To reduce the influence of large peaks, data were log2-transformed. Both quantitative (terpenoid concentrations) and qualitative (presence/absence) variation were analyzed using Manhattan and Jaccard distances, respectively, calculated with the *vegdist* function (`vegan` package). Pairwise comparisons of ancestry groups were conducted with the *pairwise.adonis* wrapper (*pairwiseAdonis* package; Martinez Arbizu, 2020), using 999 permutations for significance testing. The *betadisper* function was used to calculate Euclidean distances to group spatial medians with principal coordinate analysis. Variance differences were tested with ANOVA, followed by Tukey’s post-hoc comparisons, and patterns were visualized by plotting the first two principal coordinate axes.

## 4. Results

### 4.1 Phylogenetic analyses

Assembly and filtering with `iPyRAD` yielded 66,280 SNPs for phylogenetic analysis. On average, 19,858 loci were recovered per sample, with over half present in only four or five individuals. All ingroup and outgroup species were resolved as monophyletic with 100% bootstrap support. The ingroup (*J. grandis*, *J. occidentalis*, and *J. osteosperma*) and the broader serrate leaf juniper clade (ingroup species + *J. arizonica*, *J. californica*, and *J. deppeana*) were each monophyletic. Consistent with Uckele et al. (2021), *J. osteosperma* was basal to sister species *J. grandis* and *J. occidentalis* (Fig. 1).

### 4.2 Genetic variation across populations, species, and hybrids

Sequencing generated 930.7 million raw reads, reduced to 738.6 million reads after filtering. The mean number of assembled reads per individual was 359,245. After stringent filtering, 8,319 SNPs were retained for all samples and 9,125 SNPs for the subset of parental and hybrid (ingroup) individuals.

The PCA of all species (Fig. 2a) showed clear separation between ingroup and outgroup taxa along PC1. PC2 distinguished the three major outgroup lineages that were resolved by the phylogenetic analysis (Fig. 1): 1) *J. scopulorum*, 2) *J. californica*, and 3) *J. arizonica*/*J. deppeana*. For parental and hybrid samples, PC1 separated the three parental species, with *J. grandis* and *J. occidentalis* closer on PC1 but differentiated along PC2 (Fig. 2b). Hybrids formed a diffuse cluster between parental species clusters. PCAs within each parental species explained less variance and showed limited clustering (Fig. 2c-e). Geographic and genetic distances were not significantly correlated in *J. osteosperma* (r = -0.54, p-value = 0.89) or *J. occidentalis* (r = 0.55, p-value = 0.2), suggesting a lack of isolation by distance. Geographic distances between populations ranged from 72–859 km in *J. osteosperma*, 58–250 km in *J. occidentalis*, and 20–77 km in *J. grandis* (Fig. 2f-h).

**Figure 2.**
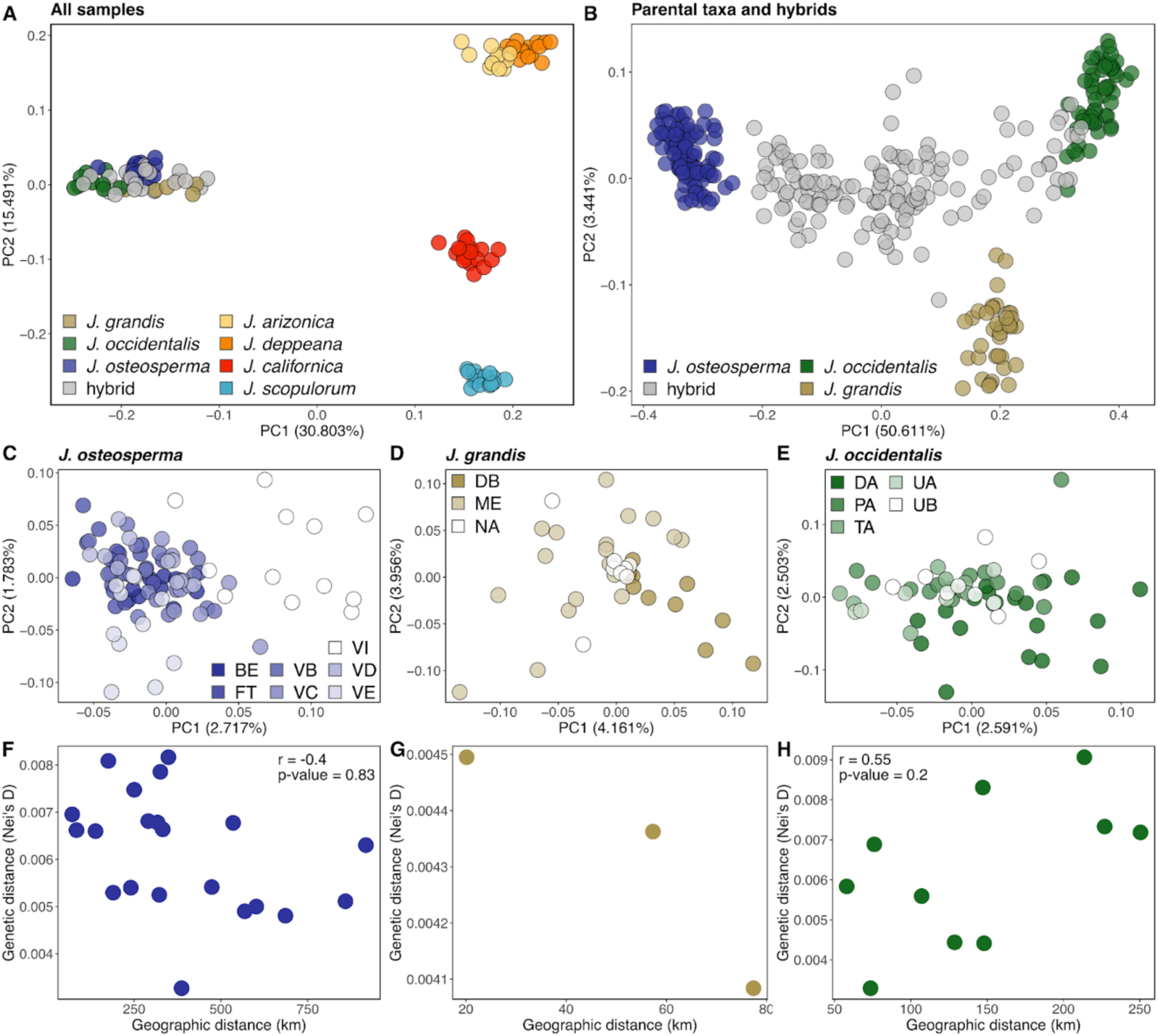
Genetic variation is strongly structured by species and hybrid status, yet intraspecific differentiation among populations is minimal. Genotype probabilities estimated with `entropy` were analyzed via PCA for (A) ingroup and outgroup individuals, and (B) parental species and hybrids, with points colored by species. Separate PCAs were then performed for each parental species (C–E), with points shaded by population. Finally, geographic vs. genetic distance relationships are shown for each parental species (F–H). No significant relationship was found for *J. osteosperma* and *J. occidentalis*, and *J. grandis* could not be tested due to limited sampling localities.

Interspecific F_ST_ values were ∼4.5 times higher on average (mean F_ST_ = 0.091) than intraspecific values (mean F_ST_ = 0.02) (Fig. 3). Genetic diversity measures (Θπ and Θw) were generally high and correlated with one another across populations (r = 0.89). Hybrids exhibited higher diversity than parental species (Supplementary Table 2–3). Tajima’s *D* was negative for all populations except UA, suggesting an excess of rare alleles consistent with population expansion (Supplementary Table 3).

**Figure 3.**
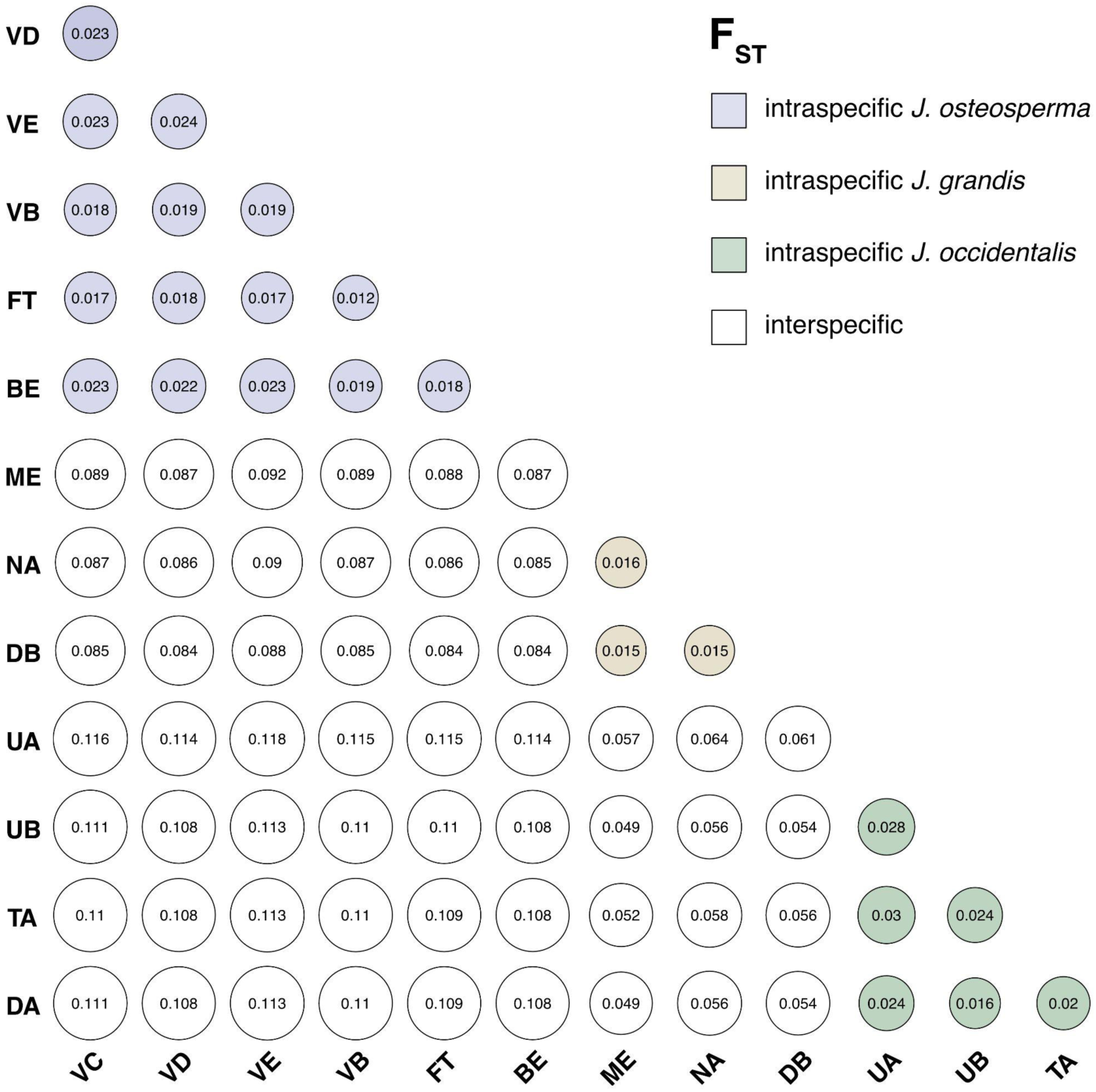
Genome-wide F_ST_ estimates are depicted in a bubble plot, where bubble size corresponds to the F_ST_ value. All pairwise estimates among parental populations are shown; additional population details can be found in Figure 1 and Supplementary Table 1. Unfilled bubbles represent interspecific F_ST_ estimates, while filled bubbles indicate intraspecific F_ST_.

A neighbor-joining tree based on Nei’s *D* mirrored the phylogeny and PCA, resolving hybrids as an evolutionary grade between *J. osteosperma* and the *J. grandis*-*J. occidentalis* clade, except for population ZA, which grouped between *J. grandis* and *J. occidentalis* (Fig. 4a). The *K* = 3 Bayesian clustering model identified the three parental species as ancestral populations. As reported previously (Adams, 2013a,b), parental species were rare within the hybrid zone. Most hybrids exhibited ancestry from all three species, but hybrid population ZA displayed primarily *J. grandis* and *J. occidentalis* ancestry (Fig. 4a). Ancestry proportions varied geographically across the hybrid zone: western hybrids possessed more equal proportions of parental ancestry, *J. osteosperma* ancestry increased with increasing longitude, and northern and southern hybrids showed higher *J. occidentalis* or *J. grandis* ancestry, respectively (Fig. 4b). In the *K* = 2 Bayesian clustering model, sister species *J. occidentalis* and *J. grandis* (hereafter referred to as the western parentals) were identified as a single source population and *J. osteosperma* was identified as the other source population. Plotting admixture proportions (*q*) against interspecific ancestry (*Q*) revealed a predominance of F_1_ and backcross hybrids, with few late-generation hybrids (3%, Fig. 4c). 37% of hybrids were backcrossed to *J. osteosperma*, 25% to the western parentals, and 24% were F_1_ hybrids, while 11% were *J. grandis* x *J. occidentalis* hybrids. A longitudinal cline in admixture proportions showed a sharp transition to admixed ancestry between 123°W and 119.5°W, followed by a more gradual shift toward *J. osteosperma* ancestry eastward of ∼119.5°W. Hybrid ancestry was most variable at the inflection point (∼119.5°W), where F_1_, western parental backcrosses, and late-generation hybrids occurred across a range of latitudes. East of this inflection point, backcross hybrids to *J. osteosperma* occur across a larger range of longitudes, extending into central Nevada (Fig. 4d).

**Figure 4.**
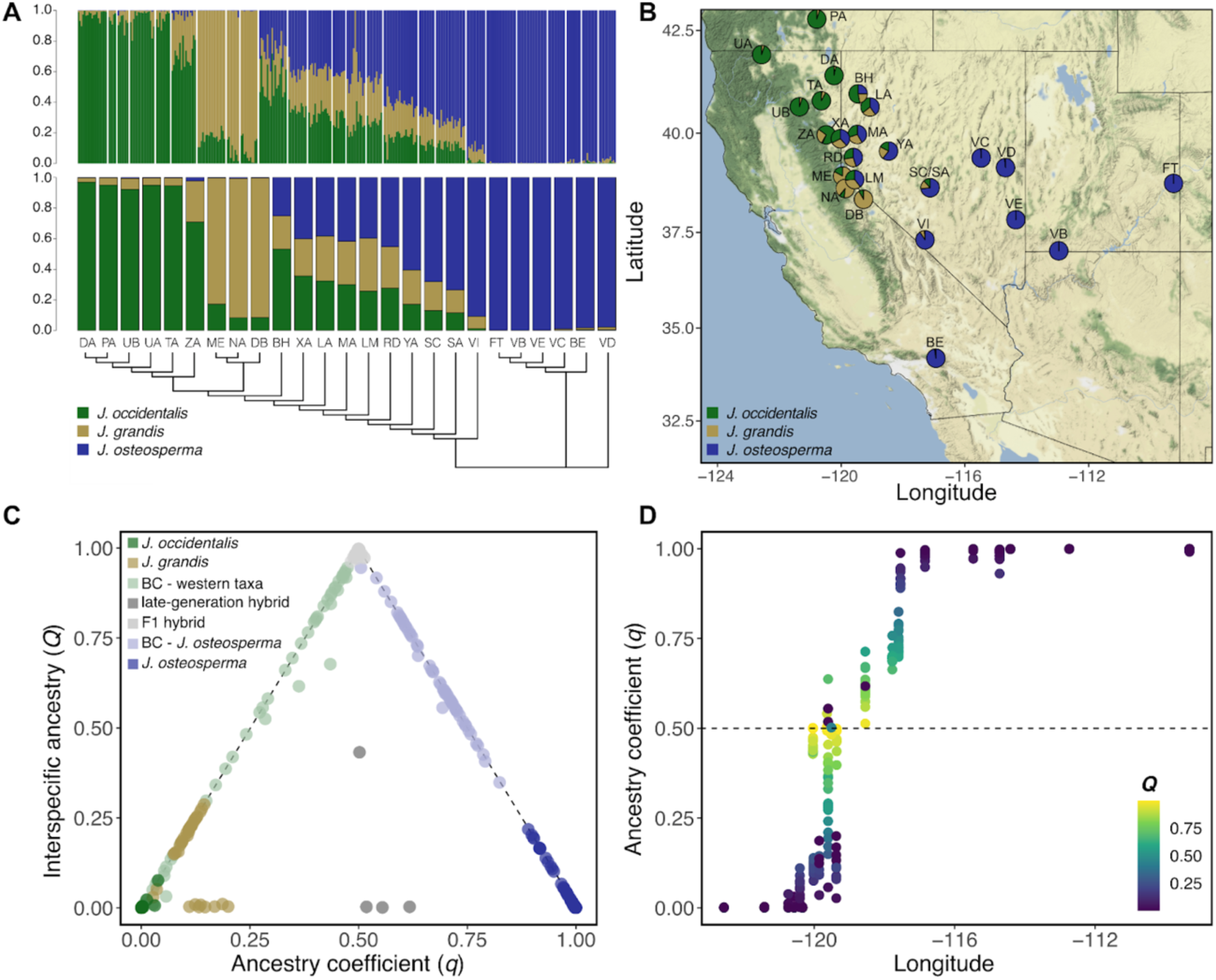
(A) The admixture model of `entropy` was set with three ancestry sources (*K* = 3), resulting in parental populations with complete or nearly complete ancestry from one of the three lineages (color-coded) and hybrid populations with mixed ancestry from two or three lineages. Individual ancestries (*q*) are shown in the upper barplot, while population mean ancestries are depicted in the lower barplot. Below the barplots, a neighbor-joining tree based on Nei’s *D* genetic distances illustrates the evolutionary relationships among populations. (B) Population mean ancestries for each sampling locality are also depicted as pie charts overlaid on a geographic map. (C) The ancestry complement model of `entropy` was set with two ancestry sources (*K* = 2) for ease of interpretation, treating sister taxa *J. grandis* and *J. occidentalis* as a single ancestral source population (see Methods). This model estimates interspecific ancestry (*Q*), the proportion of an individual’s genome with alleles inherited from different source populations. Interspecific ancestries (*Q*) are plotted against individual ancestries (*q*) for each parental and hybrid individual to distiguish hybrid classes (e.g., backcrosses, F_1_, F_2_, F_N_). (D) Individual ancestries (*q*) are plotted against longitude, with a color ramp indicating the corresponding interspecific ancestries (*Q*).

### 4.3 Ecological divergence among parental taxa and hybrids

Climate variables were highly correlated with one another (Supplementary Tables 1, 2). The DAPC of range-wide climate data revealed distinct contemporary climates for the parental species, with hybrids occupying intermediate climatic environments (Fig. 5b). Key variables differentiating species included monsoonality, minimum summer temperature, precipitation seasonality, and the fraction of actual evapotranspiration derived from monthly precipitation rather than from soil water (pdAET-SWB) (Table 1).

**Figure 5.**
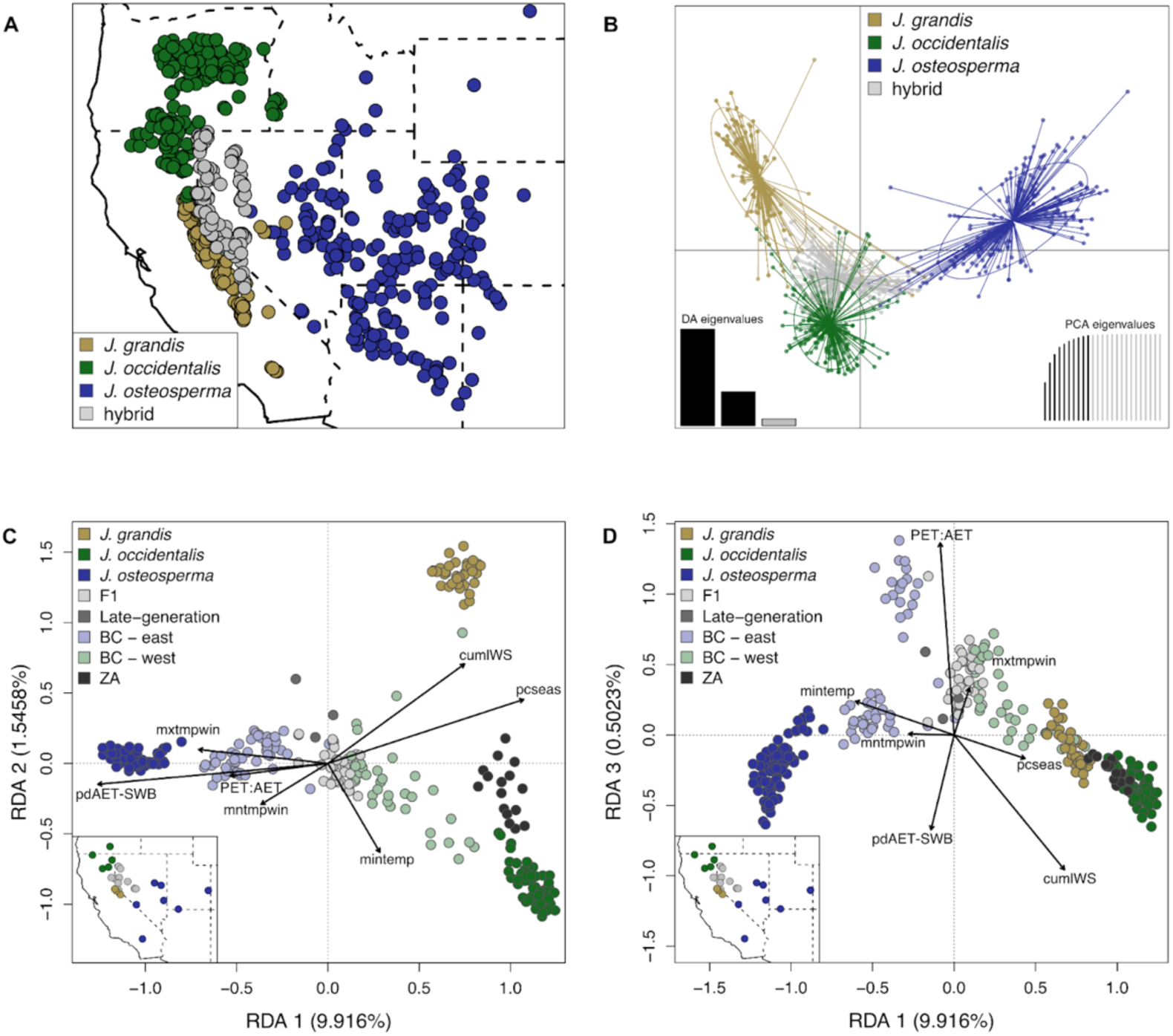
Climate variation differs among parental species and hybrids, with hybrid populations occupying environments intermediate to the parentals. (A) Two hundred geographic coordinates were randomly sampled across the ranges of each parental species and the hybrid zone. All *Juniperus osteosperma* observations west of -118° longitude were classified as hybrids based on population genetic analyses (Figure 4). (B) Climate data for each locality in panel A were analyzed using discriminant analysis of principal components (DAPC) with the first ten principal components. (C and D) Redundancy analyses (RDA) were conducted to illustrate how climate variables explain genetic variation across the hybrid zone. Unlike the DAPC in panel B, these RDAs used genotype probabilities and climate data from the 25 sampled populations (map insets, lower left). In the RDA biplots, the proximity of points (representing individuals) indicates similar genetic responses to climate predictors. Right-angle projections of points onto vectors represent variable values: smaller projections indicate larger values for a given variable, and vice versa. RDA axes 1 and 2 are shown in panel C, and axes 1 and 3 in panel D. Descriptions of climate variables are provided in Table 1.

**Table 1.** Climate variables used in various analyses. Discriminant analysis of principal components (DAPC) was performed on range-wide climate data from 200 populations across the hybrid zone and parental ranges. In contrast, redundancy analyses (RDA) and random forest models (RF_a_ and RF_b_) utilized data from the parental and hybrid populations (Figure 1). The table displays variable contributions for DAPC and variable importance for RF_a_ and RF_b_, with the five most influential variables for each model highlighted in bold. Asterisks in the first column denote variables used as predictors in the RDA model (Fig. 5c–d). RF_a_ employed a random forest classification approach to predict ancestry class, while RF_b_ used a random forest regression approach to predict hybrid admixture proportions. Variables not included in certain models are indicated as NA.

| Variable | Definition | DAPC | RF<sub>a</sub> | RF<sub>b</sub> |
| --- | --- | --- | --- | --- |
| cumlPET | Cumulative PET; amount of water that would be evapo-transpired if enough water were available | 0.0075 | **5.6** | 0.33 |
| cumlAET | Cumulative AET; actual amount of water that is evapo-transpired; proxy for productivity | 0.0195 | 2.29 | 0.11 |
| cumlSWB | Cumulative SWB; quantity of water stored in the soil from one month to the next | 0.0043 | 2.54 | 0.12 |
| cumlWS* | Cumulative WS | 0.0023 | 3.82 | 0.01 |
| cumlCWD | Cumulative CWD; proxy for drought stress | 0.0003 | **4.07** | 0.09 |
| SWB:AET | Ratio of SWB to AET; values > 1 indicate more stored soil water than used in AET | 0.0011 | 2.29 | **6.82** |
| WS:AET | Ratio of WS to AET; values > 1 indicate more water for soil water storage, runoff, or deep percolation than used in AET | NA | 4.07 | **7.99** |
| AET:CWD | Ratio of AET to CWD; values > 1 indicate mesic climate, values < 1 indicate xeric climate | 0.0042 | 4.83 | 0.28 |
| PET:AET* | Ratio of PET to AET; relative drought indicator; values > 1 indicate an unmet demand for water | 0.0031 | 3.56 | 0.02 |
| pdAET-SWB* | Positive difference between AET and SWB; fraction of AET derived from monthly precipitation rather than soil water | **0.1246** | **7.12** | 0.1 |
| pdWS-AETorSWB | Positive difference between WS and the greater of AET or SWB; cumulative water available for runoff or deep percolation | 0.0134 | 1.27 | **21.32** |
| WS:AETorSWB | Spring ratio of WS and the greater of AET or SWB; values > 1 indicate spring water available for runoff or deep percolation | 0.0031 | 3.56 | 0.27 |
| monsoon | Monsoonality; pattern of pronounced precipitation during summer months | **0.2903** | **4.83** | **10.64** |
| pcseas | Precipitation seasonality | **0.1317** | 3.05 | 0.24 |
| prcpann | Annual precipitation | 0.0009 | 2.04 | 0.003 |
| prcpspr | Spring precipitation | 0.0029 | 2.04 | 1.05 |
| prcpsum | Summer precipitation | NA | **7.38** | 1.02 |
| prcpfal | Fall precipitation | 0.0053 | 1.27 | 0.07 |
| prcpwin | Winter precipitation | NA | 2.04 | 0.34 |
| mintemp | Minimum temperature | **0.0635** | 6.87 | 0.16 |
| maxtemp | Maximum temperature | 0.0048 | 3.82 | 0.006 |
| temprang | Temperature range | 0.0539 | 2.8 | 0.05 |
| mntmpspr | Minimum spring temperature | 0.0185 | 2.54 | 0.09 |
| mxtmpspr | Maximum spring temperature | NA | 1.53 | 0.71 |
| mntmpsum | Minimum summer temperature | **0.1283** | 1.78 | 0.1 |
| mxtmpsum | Maximum summer temperature | NA | 2.04 | 0.36 |
| mntmpfal | Minimum fall temperature | 0.0355 | 3.56 | 0.01 |
| mxtmpfal | Maximum fall temperature | 0.0034 | 2.04 | 1.64 |
| mntmpwin* | Minimum winter temperature | 0.0435 | 3.31 | 0.11 |
| mxtmpwin* | Maximum winter temperature | 0.0338 | 2.54 | 0.61 |
| latitude |  | NA | NA | 0.94 |
| longitude |  | NA | NA | **6.74** |

*Note.* AET = actual evapotranspiration; PET = potential evapotranspiration; SWB = soil water
balance; WS = water supply; CWD = climate water deficit.

The random forest classification model accurately predicted ancestry class (95% CI: 96.5–100%) using climate data alone. The five most important variables, ranked by mean decrease in accuracy, were summer precipitation, pdAET-SWB, cumulative potential evapotranspiration, monsoonality, and cumulative climate water deficit (Table 1). A second random forest regression model was developed to predict hybrid ancestry proportions using climate and geographic variables (latitude and longitude). The model achieved a test set MAE of 0.04 and RMSE of 0.05, consistent with cross-validated MAE (0.05) and RMSE (0.07). The R^2^ values for cross-validation (0.77) and the test set (0.88) indicate strong predictive performance and generalization. Variable importance analysis highlighted cumulative water available for runoff or deep percolation (pdWS-AETorSWB) (21.3%) as the most influential predictor (Table 1).

The first three axes of RDA with environment and geography as predictors and genetic variation as response explained appreciable portions of variance (Fig. 5). The first two RDA axes separated the parental species into distinct clusters, with hybrids positioned between them. *Juniperus grandis* and *J. occidentalis* were associated with higher values of cumulative water supply and precipitation seasonality, while *J. osteosperma* was linked to higher values of pdAET-SWB, which was strongly correlated with summer precipitation (r = 0.71) and monsoonality (r = 0.79) (Fig. 5c). The third RDA axis separated hybrids from parental individuals, with hybrids associated with higher values of unmet demand for water and maximum winter temperature (Fig. 5d).

### 4.4 Phytochemical variation across species and hybrids

GC-MS analysis identified 163 terpenoid compounds, 55 of which were matched to known molecules, including one aliphatic, one phenylpropanoid, 15 monoterpenes, 19 diterpenes (2 bicyclic, 17 tricyclic), and 19 sesquiterpenes (6 monocyclic, 11 bicyclic, and 2 tricyclic) (Supplementary Table 4). Hybrids produced no novel terpenoids but did synthesize all 163, whereas each parental species produced only a subset (Supplementary Table 3). Hybrids differed significantly from one or more parental groups for 96 compounds (59%), with 68 (71%) classified as dominant, 14 (15%) as intermediate, 14 (15%) as transgressively high, and none as transgressively low. Among the dominant compounds, 17 (25%) were dominant toward *J. grandis*, 17 (25%) were dominant toward both *J. grandis* and *J. occidentalis*, 12 (18%) were dominant toward both *J. grandis* and *J. osteosperma*, 11 (16%) were dominant toward *J. osteosperma*, 9 (13%) were dominant toward both *J. osteosperma* and *J. occidentalis*, and 2 (3%) were dominant toward *J. occidentalis*. At least one hybrid was transgressive for 99 (61%) compounds, and total terpenoid concentrations were significantly higher in hybrids (Supplementary Fig. 4).

PCA of terpenoid data separated ancestry groups along PC1 (12.7% variance) and PC2 (8.2%), driven largely by tricyclic diterpenes (Fig. 6; Supplementary Table 4). Hybrids were differentiated from the parental groups along PC1, and intermediate to *J. osteosperma* and *J. occidentalis* along PC2 (Fig. 6a). The random forest model predicting ancestry class based on terpenoid data was highly accurate (95% CI: 85.3 - 97). RDA-based variance partitioning revealed that genetics, geography, and environment explained 11.9% of terpenoid variation, with genetic variation contributing the most (3.3%), followed by geography (2.8%) and environment (1.3%). The remaining 4.6% was confounded, indicating that the effects of genetics, geography, and climate could not be fully separated (Table 2).

**Figure 6.**
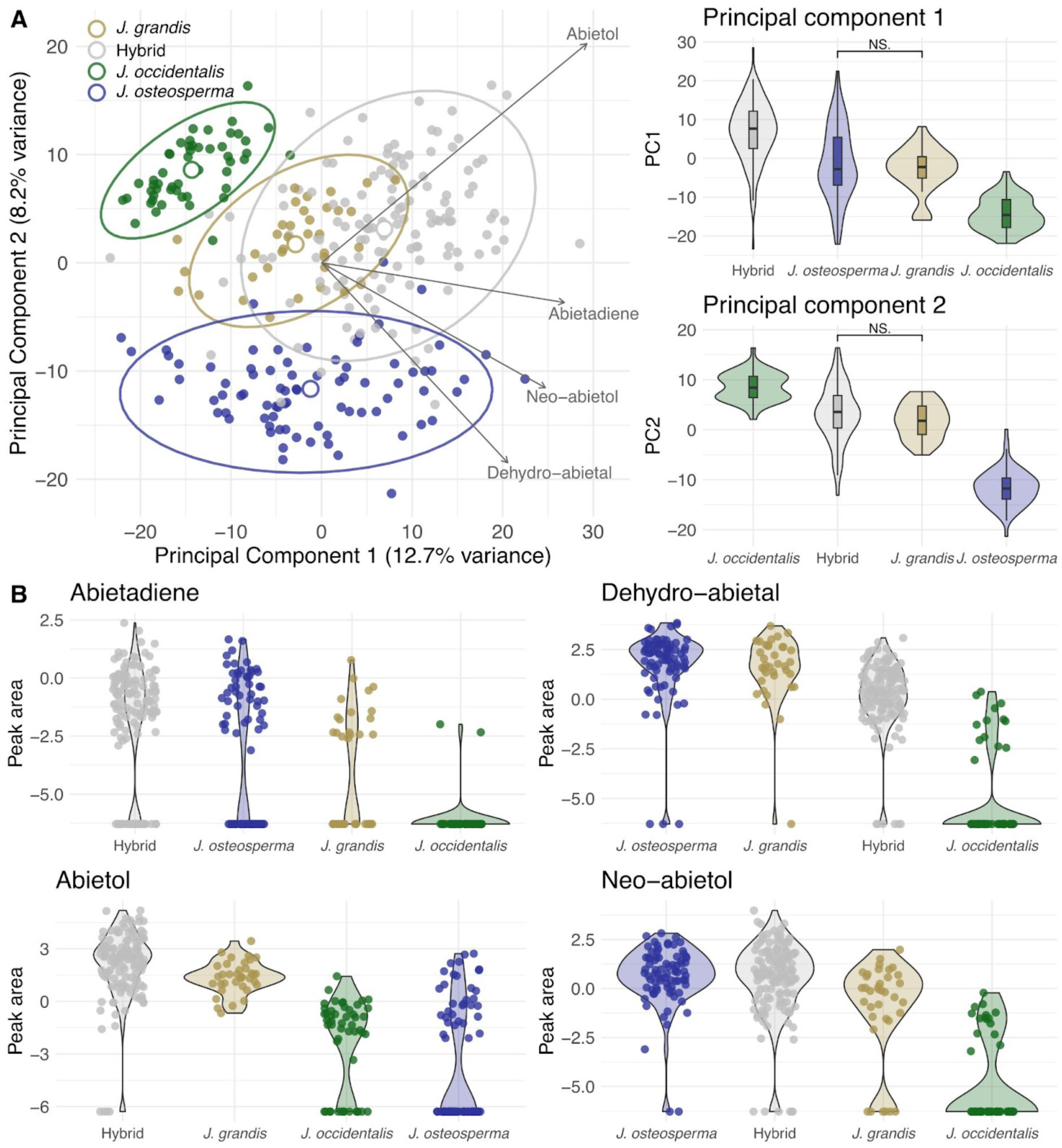
(A) Principal component analysis (PCA) of terpenoid profiles for *Juniperus* species and hybrids. Each point represents an individual sample, color-coded by ancestry group. Colored ellipses represent the 95% confidence intervals for each group, and larger open points indicate group centroids. Gray arrows illustrate the loadings of key terpenoid compounds highlighted in panel B and the text. To the right of the PCA, violin plots display the distributions of PC1 and PC2 scores for each ancestry group, ordered by their mean PC scores from highest to lowest. All pairwise comparisons of group means are significant except those indicated with brackets. (B) Violin plots showing the distributions of log_2_-transformed peak areas for four key abietane diterpenoids that significantly load onto PC1 and PC2. Jittered points within the violins represent individual sample values, highlighting variation in compound concentrations across species and hybrid groups.

**Table 2.**
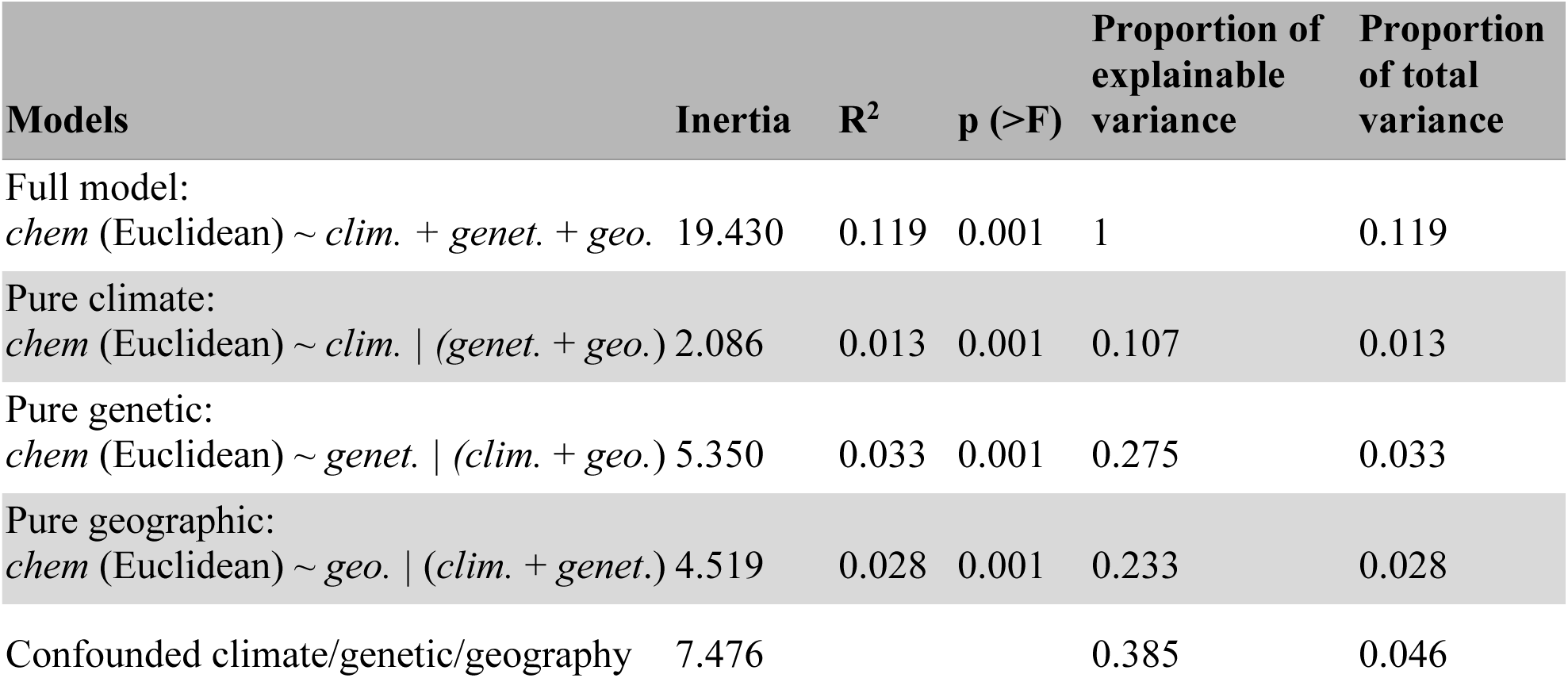
Variance partitioning results from redundancy analysis (RDA) and partial redundancy analysis (pRDA), elucidating the contributions of genetic, geographic, and environmental factors to phytochemical variation.

Alpha and gamma diversity, whether based on concentrations or presence/absence data, was highest in hybrids (Supplementary Table 5, Supplementary Fig. 5). The results of the additive beta diversity analyses are presented in the Supplementary Materials (Supplementary Results, Supplementary Fig. 6). We also used distance-based approaches to assess differences in terpenoid variability and composition across ancestry groups. The hybrid group showed the highest variance in terpenoid concentrations, while the *J. occidentalis* group exhibited the most variance in terpenoid presence/absence. PCoA of terpenoid data separated ancestry groups along the first three principal coordinates axes (Supplementary Fig. 7). Additional results from the distance-based analyses are presented in the Supplementary Materials.

## 5. Discussion

We leveraged genetic and phytochemical data to elucidate patterns of hybridization and its consequences in a group of ecologically significant *Juniperus* species of the arid west, revealing clear genetic differentiation among the parental species despite extensive hybridization in an environmentally intermediate secondary contact zone. Environmental factors predicted both hybrid occurrence and ancestry variation, suggesting that local adaptation serves as a barrier to introgression and that climate helps determine admixture gradients. An abundance of backcrosses, and a paucity of late generation hybrids, indicates that gene flow across species boundaries is a factor further shaping genetic variation in this group despite the maintenance of divergence among parental species. This syngameon structure (Boecklen 2017) not only connects and expands the environments and distributions where these species occur but also likely buffers their evolutionary potential amid rapid environmental change. By coupling landscape genomic analyses with untargeted metabolomic data, we further characterized distinct phytochemical profiles between parental species and hybrids; notably, hybrids exhibited novel variation, including increased chemical diversity and transgressive inheritance patterns for key defensive specialized metabolites, illustrating how hybridization can generate new genetic and functional variation with extended ecological consequences.

### 5.1 Genetic variation across populations, species, and hybrids

Hybridization significantly shapes evolutionary histories and often complicates phylogenetic reconstructions in plants (Rieseberg and Soltis 1991; Folk et al. 2017). In *Juniperus*, phylogenetic analyses have been particularly challenging due to reticulation. For instance, chloroplast DNA analyses (Adams et al. 2006; Adams and Schwarzbach 2013) positioned *J. grandis* and *J. osteosperma* as sister species, conflicting with morphological evidence supporting a sister relationship between *J. grandis* and *J. occidentalis*. In contrast, our analysis of thousands of nuclear loci resolved these relationships with 100% bootstrap support, confirming *J. grandis* and *J. occidentalis* as sister species (Fig. 1; see also Uckele et al. 2021).

Forest trees commonly exhibit low genetic differentiation across broad geographic ranges, even over vast distances (Petit and Hampe 2006; Neale 2007). Populations within each of the parental *Juniperus* species displayed low population genetic differentiation and no isolation by distance (Fig. 2f-h), even among *J. osteosperma* populations separated by nearly 1,000 km (Fig. 2f). Despite this, clear genetic differentiation among species is maintained, indicating reproductive barriers (Fig. 3). Similar to other North American conifers (Menon et al. 2018; Acosta et al. 2019; Haselhorst et al. 2019), genetic diversity was high within parental and hybrid populations (Supplementary Table 2). Standing genetic variation, a key predictor of adaptive potential, may be particularly important for long-lived, sessile trees facing climate change (Barnosky 1987; De Carvalho et al. 2010). Hybrids and *J. osteosperma* showed the highest genetic diversity (Supplementary Table 2), suggesting roles for admixture and large population sizes, respectively, in maintaining diversity.

*Juniperus osteosperma* occupies the largest geographic and elevational range of the three parental species. Fossil evidence indicates that during the last glacial maximum, *J. osteosperma* maintained a stable range in the Great Basin Desert and expanded southward into the Mojave Desert, southern Sierras, Central Valley, and Sonoran Desert (Thompson 1990; Nowak et al. 1994). In contrast, *J. grandis* and *J. occidentalis* reached their current ranges in the early Holocene, experiencing greater displacement and range contraction due to ice-age climate shifts (Cole 1983; Nowak et al. 1994). Juniper woodlands declined until the late Holocene, when wetter conditions during the Neoglaciation period spurred population growth and downslope expansion (Wigand et al. 1995). This history may explain the negative genome-wide Tajima’s *D* values observed in nearly all populations, with significantly lower values in *J. osteosperma* (Table 3), suggesting recent population expansion.

### 5.2 Patterns of admixture in a secondary contact zone

Contemporary hybrid zones and evidence of ancient reticulation suggest admixture has played an important role in *Juniperus* (Adams 1994, 2016; Adams et al. 2016, 2017, 2018, 2019, 2020; Farhat et al. 2020). Our study expands this understanding by documenting ancestry variation across a hybrid zone formed by admixture among three closely related species. These species underwent recurrent cycles of allopatry and secondary contact during the Quaternary, with the latest secondary contact likely beginning after the last glacial retreat in the early Holocene (∼11,700 years ago), as milder climates enabled downslope expansion (Cole 1983; Nowak et al. 1994).

The hybrid zone lacked late-generation hybrids (e.g., F_2_s, F_N_s, etc.) despite the prevalence of F_1_s, suggesting strongly asymmetric gene flow from parental populations (Barton and Hewitt, 1985) or some degree of hybrid breakdown (Burke and Arnold, 2001). An abundance of backcrosses indicates permeable but strongly maintained species boundaries among the three *Juniperus* species, typical of a syngameon, where networks of closely related species hybridize with consequences for the evolutionary dynamics of the group. Syngameons have been observed in many woody plants, including pinyon pines (Buck et al. 2023), oaks (Cronk and Suarez-Gonzalez 2018), poplars (Chhatre et al. 2018), birches (Touchette et al. 2024), and *Eucalyptus* spp. (Flores-Rentería et al. 2017). These networks of admixture may enhance resilience to environmental change by increasing effective population sizes, maintaining genetic diversity, and facilitating adaptive introgression (Cannon and Petit 2020; Buck et al. 2023).

Our results provide evidence for bidirectional backcrossing, with gene flow into the western parental genomic background confined to the California-Nevada border, while gene flow into *J. osteosperma* extends east into central Nevada (Fig. 4b,d). This pattern mirrors dominant, west-to-east wind patterns in western Nevada (the westerlies), which can shape genetic structure in wind-pollinated trees (Kling and Ackerly 2021). The transition from western parental to *J. osteosperma* ancestry forms a longitudinal cline along the ecotone between the Cascade-Sierra ranges and the Great Basin Desert (Fig. 4d). Hybridization often occurs in ecotones, where transitional zones provide contact opportunities for diverged ecotypes and reduced parental fitness favors hybrids (Barton and Hewitt 1985; Harrison 1993; Abbott 2017). Consistent with this pattern, our results suggest that the juniper parental species occupy distinct climatic zones, while hybrids are found in intermediate climates. Random forest analyses identified climate variables, particularly cumulative water available for runoff or deep percolation, as stronger predictors of hybrid ancestry proportions than geography. Together, these results indicate that adaptation to local environments is a key driver of reproductive isolation and ancestry variation across the hybrid zone, aligning with findings from other tree hybrid systems (Cullingham et al. 2012; Hamilton et al. 2013; De La Torre et al. 2014; Menon et al. 2018).

### 5.3 Phytochemical variation across species and hybrids

Known adaptive functions of plant specialized metabolites include defense, signalling, and protection from environmental stress (Pichersky and Raguso 2016). In conifers, terpenoids are central to these roles, forming the primary constituents of oleoresin, a viscous resin found throughout the plant that contains mixtures of monoterpenes, sesquiterpenes, and diterpenoid resin acids. The exudation and hardening of oleoresin in response to wounding, as well as its antimicrobial and anti-herbivore properties, make it a primary defense in conifers (Celedon and Bohlmann 2019). Here we quantified foliar terpenoid variation across a conifer hybrid zone encompassing a large range of genetic and environmental variation. Genetic variation explained the most variance in terpenoid concentrations (3.3%), followed by geography (2.8%) and environment (1.3%). The high accuracy of the random forest model and clustering by ancestry in the PCA revealed distinct chemotypes for each ancestry group, highlighting a strong genetic basis for terpenoid biosynthesis, with additional unmeasured influences that may be attributed to phenology, ontogeny, and biotic and abiotic factors (Moore et al. 2014).

Variation in terpenoid profiles among species and hybrids arises from differences in presence/absence of individual compounds (qualitative) and concentration of those compounds (quantitative). Our results show that both contribute to the unique profiles observed. While hybrids did not express novel terpenoids, their profiles were more chemically rich than those of the parental species. This richness resulted from complementarity, with hybrids expressing the full spectrum of parental terpenoids, while each parent expressed a subset. Such patterns are common in plant hybrids and align with Mendelian inheritance with dominance (Orians 2000). Hybrids were not only more chemically rich but also more variable in their terpenoid concentrations. Higher variation, often documented in plant hybrids, is likely driven by additive inheritance of compound concentrations and a broad spectrum of hybrid ancestries from backcrossing (Cheng et al. 2011). For most compounds, at least one hybrid exceeded the parental range, and 8.6% of compounds were transgressively high, with mean concentrations significantly exceeding those of the parents. Transgressive variation in hybrids generates novel phenotypic variation that can drive adaptive divergence, hybrid lineage establishment, and diversification (Lexer et al. 2003; Buerkle and Rieseberg 2008; Kagawa and Takimoto 2018). While studies show that hybrids are generally as susceptible or more to herbivores than their parents, some show greater resistance, offering novel adaptive variation that may spread through introgression (Cheng et al. 2011). In juniper hybrids, the transgressive expression of two biosynthetic precursors of abietic acid, abietadiene and abietol (Fig. 6b), suggests adaptive potential. Abietic acid, a diterpene resin acid, is central to conifer defense (Keeling & Bohlmann 2006; Oh et al. 2017). Although we could not directly assess abietic acid expression in the hybrids, the transgressive expression of its precursors suggests the potential for transgressive expression of abietic acid as well. Quantifying resin acids in natural juniper hybrids and assessing herbivore resistance could clarify the adaptive significance of transgressive phytochemical variation in this hybrid zone.

Although phytochemical variation in hybrids is well-documented (Rieseberg et al. 1993; Orians 2000; Cheng et al. 2011; Caseys et al. 2015), its ecological and evolutionary consequences in foundational tree hybrid zones remain largely unexplored, limiting our understanding of which aspects of hybrid phytochemistry influence communities and ecosystems (Wetzel and Whitehead 2020; Volf et al. 2024). Theory and empirical studies suggest phytochemical diversity can affect herbivorous insect diversity in contrasting ways (Jones and Lawton 1991). For instance, higher phytochemical richness in Protieae species was associated with lower herbivore diversity, likely due to increased challenges for herbivores (Salazar et al. 2018). Alternatively, higher chemical diversity may facilitate host shifts or provide enemy free space, potentially increasing herbivore diversity, as observed in *Piper* species (Richards et al. 2015). However, no studies have specifically examined this relationship in hybrid zones. Although tree hybrid zones frequently host more diverse arthropod communities than parental populations (Whitham et al. 1994), the role of phytochemistry in shaping this diversity remains uncertain. Investigating patterns of arthropod diversity in this juniper hybrid zone, and its relationship to different dimensions of phytochemical diversity, could provide insights into the ecological impacts of hybridization in foundational trees.

## 6. Funding

This work was supported by the American Genetic Association EECG award to KAU, who was also supported by the National Science Foundation Graduate Research Award (Award No. 1650114). JPJ received support from the Modelscape Consortium through NSF funding (Award No. OIA-2019528).

## 7. Acknowledgements

Illumina sequencing was conducted by the Genomic Sequencing and Analysis Facility at UT Austin’s Center for Biomedical Research Support (RRID#: SCR_021713). We extend our gratitude to Oren Shelef for his assistance with population sampling and to Thomas Dilts and Sarah Barga for their support in obtaining climate data for this study. Additionally, we thank Genalynn Joy Lapira, Regina Gojar, and Brianna Jones for their help with chemical extractions.

## 8. Data Availability

We have deposited the primary data underlying these analyses in public repositories as follows:

- Environmental data, terpenoid concentration data, and ddRADseq genotype probability matrix: Available on Dryad.
- Raw DNA sequence reads: Accessible via NCBI SRA under Bioproject PRJNA1037072.

## Notes

### Competing Interest Statement

The authors have declared no competing interest.

