## Supplemental Materials for "Admixture and environment shape population genetic and phytochemical variation across a conifer hybrid zone"

**Running title:** Genetic and phytochemical variation in conifer hybrids

Kathryn A. Uckele^1^

Casey S. Philbin^2^

Lora A. Richards^2,3,4^

Lee A. Dyer^2,3,4^

Joshua P. Jahner^5,6^

Thomas L. Parchman^3,4^

^1^Division of Biological Sciences, University of Montana, Missoula, MT 59812, USA

^2^Hitchcock Center for Chemical Ecology, University of Nevada, Reno, NV 89557, USA

^3^Program in Ecology, Evolution, and Conservation Biology, University of Nevada, Reno, NV 89557, USA

^4^Department of Biology, University of Nevada, Reno, NV 89557, USA

^5^Department of Botany, University of Wyoming, Laramie, WY 82071, USA

^6^Department of Biology, New Mexico Institute of Mining and Technology, Socorro, NM 87801, USA

**Supplementary Methods**

**Ipyrad parameterization**

Briefly, reads were *de novo* assembled within individuals using vsearch (v. 2.14.1; Rognes et al., 2016) and aligned with muscle (v. 3.8.155; Edgar, 2004) to produce stacks of highly similar reads with over 85% sequence similarity (*clust_threshold*). Consensus sequences with more than 5% ambiguous bases (*max_Ns_consens*) or 5% heterozygous sites (*max_Hs_consens*) likely represent poorly aligned regions and were discarded. The consensus sequences were then clustered across individuals. Any stacks with more than 8 indels (*max_Indels_locus*), 20% variable sites (*max_SNPs_locus*), or one heterozygous site shared across more than 50% of the samples (*max_shared_Hs_locus*) are indicative of poor alignment or paralogy and were discarded. To avoid overfiltering missing data, we retained all stacks that were present in at least four samples (*min_samples_locus*).

**Gas chromatography-mass spectroscopy protocol**
Extracts (1 μL) were injected onto an Agilent 7890A gas chromatograph coupled to an Agilent 5975C quadrupole mass spectrometer (GC-MS) equipped with an Agilent HP-5MS, (5%-Phenyl)-methylpolysiloxane, 30 m x 250 µm x 0.25 µm capillary column (Agilent Technologies Inc., Santa Clara, CA, USA). Injections were split 1:10 at an inlet temperature of 250 °C, pressure of 7.1 psi, total He flow of 14 mL/min, septum purge flow of 3 mL/min, and a split flow of 10 mL/min. The 18.25 minute run began with an oven temperature of 40 °C with He carrier gas flow of 1 mL/min at 7.1 psi. This temperature was held for two minutes before elevating temperature at 20 °C/min to 325 °C and holding at 325 °C for an additional two minutes. The electron impact source (70 eV) temperature was 230 °C with the MS Quad temperature set to 150 °C. An *n*-alkane standard mix (Restek 31633, C_10_-C_40_) was injected at the start of each day for retention index calibration.

**GC-MS data processing**

Raw Chemstation GC-MS chromatograms were converted so they could be opened in MassHunter Qualitative Analysis using GC/MS Translator (Agilent Technologies Inc., Santa Clara, CA, USA). In MassHunter, total ion chromatograms (TIC) were integrated using the Agile integrator in the “find compounds by integration” feature. The resulting retention time (rt) and peak area list was tabulated using the GCalignR package (Ottensmann et al., 2018) in R using a maximum retention time (rt) difference of 0.05 min from the mean for group inclusion, and a minimum rt difference of 0.1 min between peak groups. Tabulated peak areas were normalized to n-Eicosane internal standard area and plant mass before statistical analysis.

**Additive beta diversity**

Additive beta diversity, representing the contribution of between-sample terpenoid differences to overall species terpenoid diversity, was calculated as the difference between gamma diversity and mean alpha diversity, both measured using the Shannon Diversity Index. Additive beta diversity was evaluated both qualitatively using presence/absence data (β_PA_) and quantitatively using concentration data (β_CONC_). Beta diversity can result from the replacement of compounds between samples (turnover) or from the gain and loss of compounds across samples (nestedness). To disentangle these components within ancestry groups, we converted terpenoid concentrations to presence/absence data and calculated dissimilarity indices with the betapart package in R (Baselga et al. 2018). To calculate the distribution of dissimilarity measures while accounting for uneven sampling across ancestry groups, we subsampled 24 individuals per group, repeating this procedure for 100 iterations using the *beta.sample* function.

**Supplementary Results**

**Additive beta diversity and distance-based analyses**

In the following sentences, the first number in parentheses represents additive beta diversity based on terpenoid concentrations, and the second represents that based on terpenoid presence/absence. Additive beta diversity was highest in *J. occidentalis* (β_CONC_ = 2.6, β_PA_ = 2.8) and similar across the other groups: *J. osteosperma* (β_CONC_ = 2.0, β_PA_ = 2.3), hybrid (β_CONC_ = = 2.1, β_PA_ = 2.1), and *J. grandis* (β_CONC_ = 2.1, β_PA_ = 2.2). For all ancestry groups, the turnover components of beta diversity were much higher than the nestedness components (Supplementary Fig. 6), suggesting that beta diversity was largely determined by the replacement of terpenoid compounds rather than by differences in terpenoid richness across samples.

We also used distance-based approaches to assess beta diversity within and across ancestry groups. To distinguish quantitative (concentration) from qualitative (presence/absence) variation in terpenoid profiles, we analyzed two distance matrices: Manhattan distances based on terpenoid concentrations and Jaccard distances based on terpenoid presence/absence. In the concentration-based analysis, the group variances were significantly different from one another. The hybrid group had the highest variance, followed by *J. osteosperma* and *J. grandis*, which did not differ significantly from each other, and *J. occidentalis*, which had the smallest variance. Since the group variances of *J. osteosperma* and *J. grandis* were not significantly different, we used PERMANOVA to test for differences in their terpenoid compositions and found a significant result. In the presence/absence-based analysis, *J. occidentalis* exhibited the highest variance, significantly differing from *J. grandis* and the hybrids, but not from *J. osteosperma*. This aligns with the higher additive beta diversity and greater turnover observed in *J. occidentalis* (Supplementary Fig. 6), indicating more variation in terpenoid composition within *J. occidentalis* compared to other groups. Since the group variances of *J. osteosperma*, *J. grandis*, and the hybrids were not significantly different, and the same was true for *J. occidentalis* and *J. osteosperma*, we used PERMANOVA to compare them and found significant differences in their compositions. Finally, we used PCoA to visualize patterns in the distance matrices, revealing distinct clustering by ancestry group, further supporting the differences in terpenoid profiles based on ancestry (Supplementary Fig. 7).

**Supplementary Table & Figures**

**Table 1.** Information on collection localities for focal and outgroup *Juniperus* taxa from the western United States. See Figure 1 in the main text for a geographic view of the focal localities.

| ID | Taxon label | Locality | N | Longitude (°W) | Latitude (°N) | Elevation (m) |
| --- | --- | --- | --- | --- | --- | --- |
| *Focal taxa* | |  |  |  |  |  |
| BE | *J. osteosperma* | San Bernardino, CA | 11 | 34.343 | -116.842 | 1465 |
| BH | hybrid | Buffalo Hills, NV | 18 | 40.885 | -119.604 | 1548 |
| DA | *J. occidentalis* | Parker Creek, CA | 19 | 41.467 | -120.333 | 1690 |
| DB | *J. grandis* | Fales Hot Spring Pass, CA | 11 | 38.347 | -119.371 | 2297 |
| FT | *J. osteosperma* | Fisher Towers, UT | 19 | 38.724 | -109.309 | 1443 |
| LA | hybrid | Gerlach, NV | 12 | 40.688 | -119.361 | 1189 |
| LM | hybrid | Leviathan Mine, NV | 12 | 38.774 | -119.604 | 1954 |
| MA | hybrid | south of Pyramid Lake, NV | 8 | 39.88 | -119.513 | 1171 |
| ME | *J. grandis* | Meyers, CA | 18 | 38.851 | -120.021 | 1946 |
| NA | *J. grandis* | Grover Waterfalls, CA | 8 | 38.698 | -119.857 | 1821 |
| PA | *J. occidentalis* | Paisley, OR | 16 | 42.684 | -120.572 | 1425 |
| RD | hybrid | Virginia Mountains, NV | 10 | 39.384 | -119.638 | 1792 |
| SA | hybrid | Humboldt-Toiyabe NF, NV | 19 | 38.869 | -117.596 | 2098 |
| SC | hybrid | Humboldt-Toiyabe NF, NV | 10 | 38.892 | -117.778 | 2054 |
| TA | *J. occidentalis* | north of Eagle Lake, CA | 7 | 40.824 | -120.739 | 1693 |
| UA | *J. occidentalis* | Hornbrook, CA | 5 | 41.916 | -122.573 | 687 |
| UB | *J. occidentalis* | Subway Cave Lava Tubes, CA | 9 | 40.692 | -121.420 | 1329 |
| VB | *J. osteosperma* | Coral Pink Sand Dunes, UT | 19 | 37.035 | -112.734 | 1795 |
| VC | *J. osteosperma* | Little Antelope Summit, NV | 9 | 39.397 | -115.469 | 2260 |
| VD | *J. osteosperma* | Cave Lake State Park, NV | 10 | 39.187 | -114.718 | 2186 |
| VE | *J. osteosperma* | Cathedral Gorge, NV | 11 | 37.819 | -114.412 | 1446 |
| VI | *J. osteosperma* | Lida, NV | 12 | 37.441 | -117.544 | 2134 |
| XA | hybrid | Hallelujah Junction, NV | 15 | 39.788 | -120.038 | 1561 |
| YA | hybrid | Stillwater, NV | 22 | 39.522 | -118.547 | 1187 |
| ZA | hybrid | north of Sierraville, CA | 16 | 39.642 | -120.368 | 1495 |
| *Outgroup taxa* | |  |  |  |  |  |
| GB | *J. californica* | Culp Valley, CA | 5 | 33.23 | -116.461 | 1012 |
| HA | *J. californica* | Culp Valley, CA | 10 | 33.23 | -116.461 | 1012 |
| WB | *J. scopulorum* | Rocky Mountain NP, CO | 10 | 40.514 | -105.589 | 3184 |
| WC | *J. scopulorum* | Estes Park, CO | 5 | 40.369 | -105.538 | 2356 |
| ZB | *J. deppeana* | Paradise Cemetery, AZ | 16 | 31.932 | -109.208 | 1695 |
| ZC | *J. arizonica* | Paradise Road, AZ | 14 | 31.932 | -109.205 | 1695 |

**Table 2.** Genetic diversity (𝜃_𝜋_ and 𝜃_𝑊_) and Tajima’s *D* (mean and 95% confidence interval) was calculated for each focal locality listed in Supplementary Table 1.

| Locality | Taxon label | 𝜃_𝜋_ | 𝜃_𝑊_ | Tajima’s *D* range (mean) |
| --- | --- | --- | --- | --- |
| BE | *J. osteosperma* | 0.01147 | 0.01292 | -0.43797 – -0.43151 (-0.43474) |
| BH | hybrid | 0.01184 | 0.01323 | -0.37989 – -0.37393 (-0.37691) |
| DA | *J. occidentalis* | 0.01073 | 0.01099 | -0.20873 – -0.20157 (-0.20515) |
| DB | *J. grandis* | 0.01007 | 0.01042 | -0.21415 – -0.20793 (-0.21104) |
| FT | *J. osteosperma* | 0.01132 | 0.01417 | -0.62874 – -0.62218 (-0.62546) |
| LA | hybrid | 0.01198 | 0.01394 | -0.49811 – -0.49213 (-0.49512) |
| LM | hybrid | 0.01167 | 0.01284 | -0.36621 – -0.36061 (-0.36341) |
| MA | hybrid | 0.01185 | 0.01314 | -0.40167 – -0.39627 (-0.39897) |
| ME | *J. grandis* | 0.01083 | 0.01108 | -0.19082 – -0.18404 (-0.18743) |
| NA | *J. grandis* | 0.01015 | 0.01007 | -0.10215 – -0.0966 (-0.09938) |
| PA | *J. occidentalis* | 0.01055 | 0.01087 | -0.22399 – -0.21724 (-0.22062) |
| RD | hybrid | 0.01198 | 0.01346 | -0.4246 – -0.4188 (-0.4217) |
| SA | hybrid | 0.01190 | 0.01525 | -0.67098 – -0.66486 (-0.66792) |
| SC | hybrid | 0.01187 | 0.01365 | -0.48431 – -0.47852 (-0.48142) |
| TA | *J. occidentalis* | 0.01080 | 0.01067 | -0.10837 – -0.10254 (-0.10545) |
| UA | *J. occidentalis* | 0.01082 | 0.01033 | 0.01613 – 0.02197 (0.01905) |
| UB | *J. occidentalis* | 0.01067 | 0.01041 | -0.0852 – -0.07911 (-0.08216) |
| VB | *J. osteosperma* | 0.01115 | 0.01283 | -0.45228 – -0.44551 (-0.4489) |
| VC | *J. osteosperma* | 0.01114 | 0.01273 | -0.47911 – -0.4729 (-0.47601) |
| VD | *J. osteosperma* | 0.01141 | 0.01304 | -0.48033 – -0.47407 (-0.4772) |
| VE | *J. osteosperma* | 0.01101 | 0.01203 | -0.36433 – -0.35788 (-0.3611) |
| VI | *J. osteosperma* | 0.01113 | 0.01177 | -0.27079 – -0.26408 (-0.26743) |
| XA | hybrid | 0.01104 | 0.01147 | -0.22727 – -0.22084 (-0.22406) |
| YA | hybrid | 0.01190 | 0.01459 | -0.56472 – -0.55837 (-0.56155) |
| ZA | hybrid | 0.01117 | 0.01224 | -0.35071 – -0.34433 (-0.34752) |

**Table 3.** Genetic diversity (𝜃_𝜋_ and 𝜃_𝑊_) and Tajima’s *D* (mean and 95% confidence interval) was calculated for the parental species.

| Species | 𝜃_𝜋_ | 𝜃_𝑊_ | Tajima’s *D* range (mean) |
| --- | --- | --- | --- |
| *J. osteosperma* | 0.01150 | 0.01744 | -0.90137 – -0.89458 (-0.89798) |
| *J. occidentalis* | 0.01090 | 0.01221 | -0.35958 – -0.35187 (-0.35573) |
| *J. grandis* | 0.01089 | 0.01230 | -0.37383 – -0.36692 (-0.37037) |

**Table 4.** Annotated compounds from the GC-MS analysis of hybrid and parental individuals. Of 163 peak bins, 55 were matched to known compounds and annotated. Retention time (RT), retention index (RI), the difference between the retention index of the matched compound and the observed retention index (Delta RI), and match score (Match) are provided for each annotation. Compounds identified as significant in the PCA or random forest analysis are indicated in the Analyses column.

| RT | RI | Delta RI | Match | Name | Class | Oxidation | Analyses |
| --- | --- | --- | --- | --- | --- | --- | --- |
| 5.6321 | 913 | 11 | 71 | alpha-Thujene | Monoterpene | Hydrocarbon | RF |
| 6.1126 | 970 | 1 | 93 | Sabinene | Monoterpene | Hydrocarbon | RF |
| 6.5286 | 1020 | 19 | 82 | delta-2-Carene | Monoterpene | Hydrocarbon |  |
| 6.8944 | 1064 | 62 | 85 | alpha-phellandrene | Monoterpene | Hydrocarbon | RF |
| 7.0099 | 1077 | 36 | 65 | trans-sabinene hydrate | Monoterpene | Monohydric |  |
| 7.2797 | 1110 | 26 | 64 | trans-para-menth2-en-1-ol | Monoterpene | Monohydric |  |
| 7.6792 | 1156 | 4 | 88 | Z-isocitral | Monoterpene | Aldehyde |  |
| 7.708 | 1161 | 20 | 95 | camphor | Monoterpene | Ketone |  |
| 7.7737 | 1168 | 70 | 62 | heptenol acetate | Aliphatic | Ester |  |
| 7.9012 | 1184 | 19 | 86 | Borneol | Monoterpene | Monohydric |  |
| 7.9522 | 1189 | 16 | 94 | terpinen-4-ol | Monoterpene | Monohydric |  |
| 8.1876 | 1220 | 16 | 92 | verbenone | Monoterpene | Ketone |  |
| 8.7161 | 1296 | 13 | 94 | isobornyl acetate | Monoterpene | Monohydric |  |
| 8.9444 | 1322 | 0 | 92 | methyl geranate | Monoterpene | Ester |  |
| 9.0267 | 1339 | 194 | 66 | para-menth-3-en-8-ol | Monoterpene | Monohydric |  |
| 9.854 | 1461 | 7 | 94 | pinchotene acetate | Monoterpene | Aromatic, dihydroxy |  |
| 10.197 | 1516 | 23 | 87 | trans-muurola-4(14),5-diene | Bicyclic sesquiterpene | Hydrocarbon |  |
| 10.316 | 1534 | 21 | 93 | gamma-cadinene | Bicyclic sesquiterpene | Hydrocarbon | PC2 |
| 10.382 | 1545 | 10 | 95 | elemicin | Phenylpropanoid |  |  |
| 10.501 | 1563 | 17 | 92 | hedycaryol | Monocyclic sesquiterpene | Monohydric |  |
| 10.718 | 1597 | 24 | 90 | germacrene D-4-ol | Monocyclic sesquiterpene | Monohydric |  |
| 10.784 | 1613 | 63 | 91 | cis-muurol-5-en-4-beta-ol | Bicyclic sesquiterpene | Monohydric |  |
| 10.884 | 1626 | 19 | 66 | beta-oplopenone | Bicyclic sesquiterpene | Ketone |  |
| 10.979 | 1642 | 24 | 66 | epi-cedrol | Tricyclic sesquiterpene | Monohydric |  |
| 11.023 | 1655 | 33 | 82 | 5-epi-7-epi-alpha-eudesmol | Bicyclic sesquiterpene | Monohydric |  |
| 11.087 | 1662 | 18 | 89 | alpha-muurolol | Bicyclic sesquiterpene | Monohydric | PC2, RF |
| 11.191 | 1682 | 76 | 92 | 10-epi-gamma-eudesmol | Bicyclic sesquiterpene | Monohydric |  |
| 11.235 | 1683 | 34 | 79 | beta-eudesmol | Bicyclic sesquiterpene | Monohydric |  |
| 11.353 | 1710 | 36 | 65 | analog: 8-alpha-11-elemodiol | Monocyclic sesquiterpene | Dihydroxy | RF |
| 11.629 | 1757 | 11 | 86 | 8-alpha-11-elemodiol | Monocyclic sesquiterpene | Dihydroxy | RF |
| 11.636 | 1760 | 21 | 93 | oplopanone | Bicyclic sesquiterpene | Monohydric ketone |  |
| 11.813 | 1792 | 0 | 91 | 8-alpha-acetoxyelemol | Monocyclic sesquiterpene | Dihydroxy ester |  |
| 12.007 | 1827 | 42 | 80 | Flourensadiol | Tricyclic sesquiterpene | Dihydroxy | PC1, RF |
| 12.078 | 1844 | 52 | 74 | 8-alpha-acetoxyelemol | Monocyclic sesquiterpene | Dihydroxy ester |  |
| 12.148 | 1854 | 79 | 76 | 2-alpha-hydroxy-amorpha-4,7(11)-diene | Bicyclic sesquiterpene | Monohydric |  |
| 12.365 | 1894 | 7 | 64 | Oplopanoyl acetate | Bicyclic sesquiterpene | Monohydric ketone |  |
| 13.064 | 2036 | 49 | 95 | Manool oxide | Tricyclic diterpene | Pyran |  |
| 13.32 | 2087 | 28 | 85 | 13-epi-manool | Bicyclic diterpene | Monohydric |  |
| 13.506 | 2131 | 44 | 88 | Abietadiene | Tricyclic diterpene | Hydrocarbon | PC1 |
| 14.06 | 2256 | 34 | 86 | sclareol | Bicyclic diterpene | Dihydroxy |  |
| 14.165 | 2276 | 36 | 66 | analog: abieta-7,13-dien-3-one | Tricyclic diterpene | Ketone |  |
| 14.196 | 2286 | 27 | 81 | analog: abieta-7,13-dien-3-one | Tricyclic diterpene | Ketone |  |
| 14.274 | 2298 | 32 | 71 | dehydro-abietal | Tricyclic diterpene | Aldehyde | PC2, RF |
| 14.295 | 2305 | 140 | 67 | 7-alpha-hydroxy-trans-totarol | Tricyclic diterpene | Dihydroxy |  |
| 14.46 | 2346 | 48 | 94 | 4-epi-abietal | Tricyclic diterpene | Aldehyde |  |
| 14.486 | 2358 | 46 | 99 | abieta-7,13-dien-3-one | Tricyclic diterpene | Ketone | PC1 |
| 14.687 | 2395 | 52 | 73 | analog: abietol | Tricyclic diterpene | Monohydric |  |
| 14.668 | 2396 | 5 | 71 | analog: abietol | Tricyclic diterpene | Monohydric |  |
| 14.777 | 2426 | 113 | 85 | analog: abieta-7,13-dien-3-one | Tricyclic diterpene | Ketone | PC1 |
| 14.888 | 2447 | 46 | 93 | abietol | Tricyclic diterpene | Monohydric | PC1, PC2, RF |
| 15.016 | 2486 | 0 | 61 | 3-alpha-14,15-dihydro-manool oxide | Tricyclic diterpene | Monoxy | PC1 |
| 15.057 | 2487 | 190 | 65 | analog: abieta-7,13-dien-3-one | Tricyclic diterpene | Ketone |  |
| 15.148 | 2515 | 47 | 77 | neo-abietol | Tricyclic diterpene | Monohydric | PC1 |
| 15.187 | 2523 | 34 | 62 | Hinokienone | Tricyclic diterpene | Monohydric ketone | PC2 |
| 15.406 | 2580 | 38 | 80 | Totarolone | Tricyclic diterpene | Monohydric ketone | PC1 |

*Note*. PC1 = Principal Component 1; PC2 = Principal Component 2; RF = Random Forest.

**Table 5:** Mean alpha diversity and gamma diversity, calculated with terpenoid presence/absence or concentrations, are provided for each ancestry group.

| Ancestry group | Mean alpha diversity (presence/absence) | Mean alpha diversity (concentration) | Gamma diversity (presence/absence) | Gamma diversity (concentration) |
| --- | --- | --- | --- | --- |
| Hybrid | 55.7 | 28.2 | 111.7 | 56.6 |
| *J. grandis* | 40.8 | 21.9 | 87.5 | 44.9 |
| *J. occidentalis* | 29.7 | 15.9 | 80.7 | 39.2 |
| *J. osteosperma* | 46.9 | 23.3 | 102.1 | 44.3 |

**Figure 1.** Correlations among moisture-related variables. Descriptions of each variable can be found in the main text (Table 1).

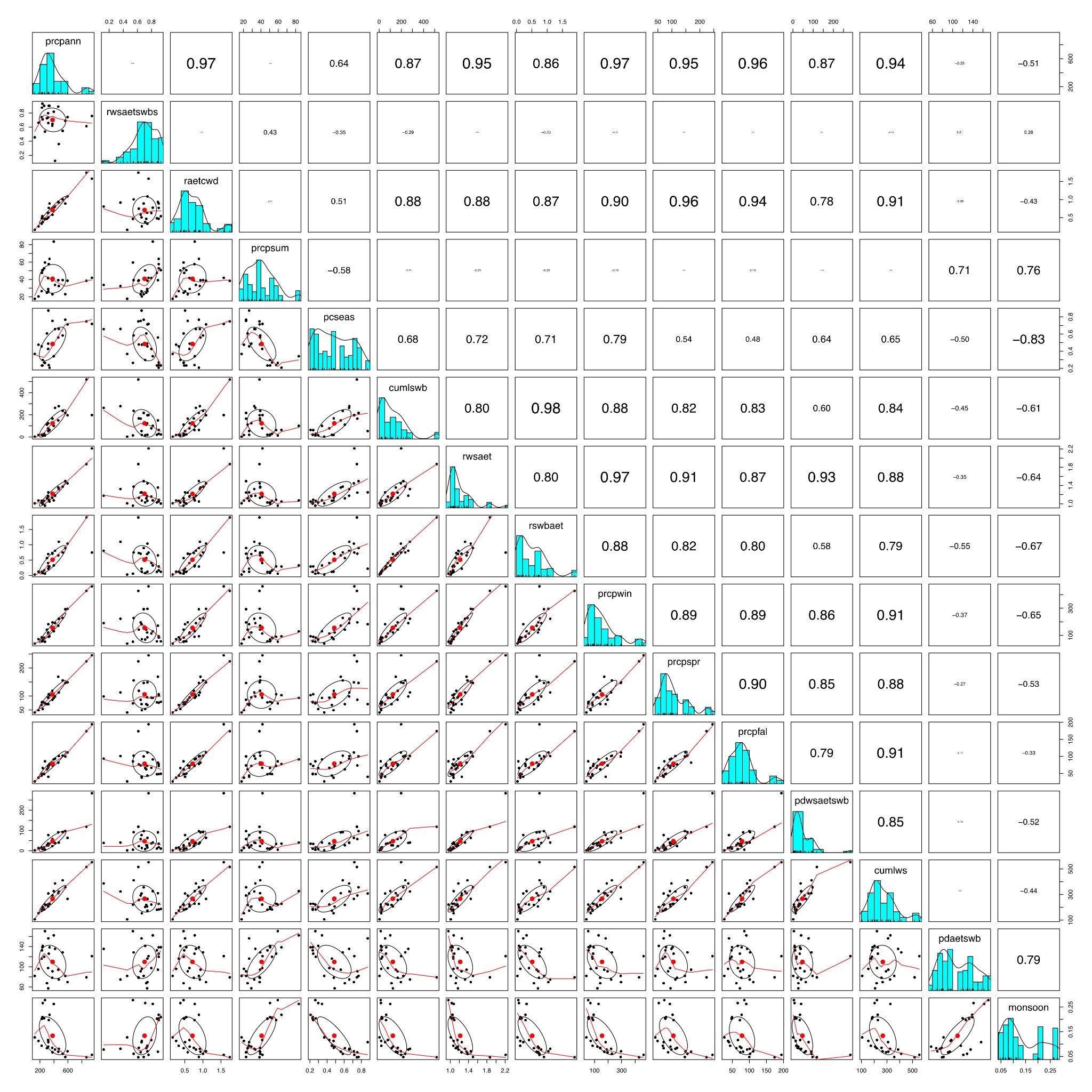

**Figure 2.** Correlations among temperature-related variables. Descriptions of each variable can be found in the main text (Table 1).

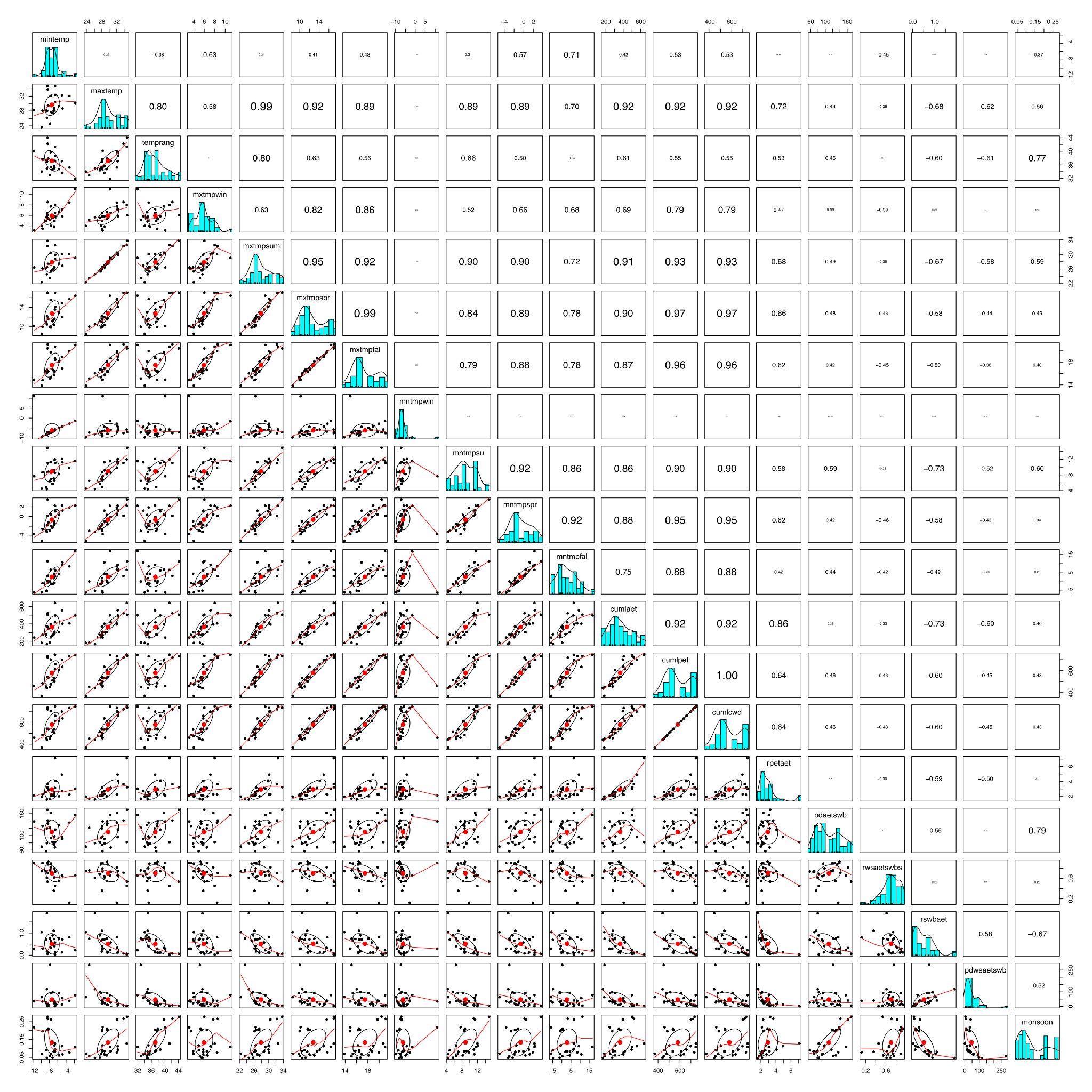

**Figure 3.** Venn diagram illustrating the overlap of 163 terpene metabolites across species and hybrid groups. Overlapping regions represent metabolites shared between two or more groups. Note that sample sizes vary across ancestry groups, which may influence the observed overlap and lack thereof.

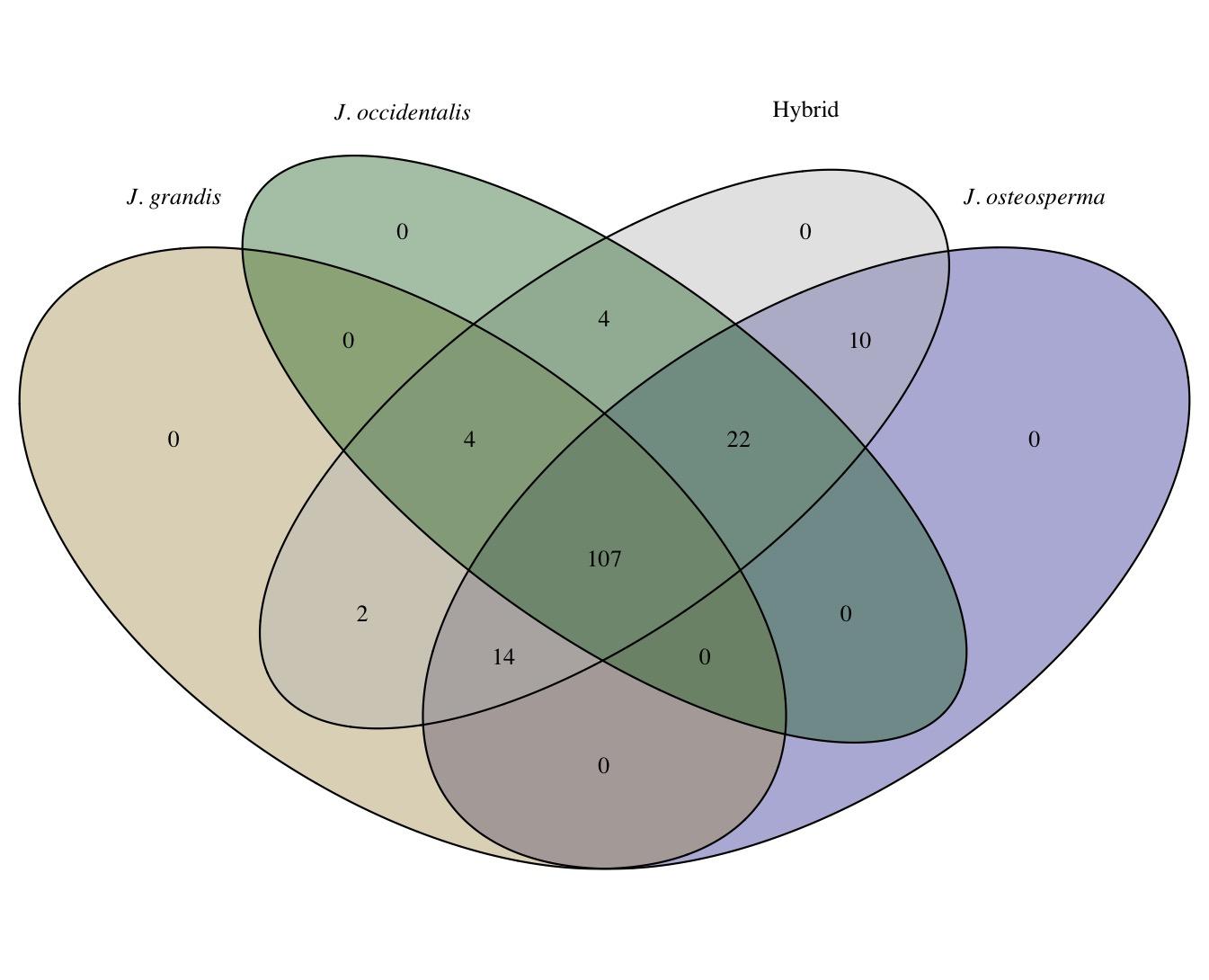

**Figure 4.** Total terpenoid concentrations summed across compounds for each ancestry group. Hybrids exhibited the highest overall terpenoid concentrations, followed by *J. osteosperma* and *J. grandis*, which were not significantly different from each other, and then *J. occidentalis*.

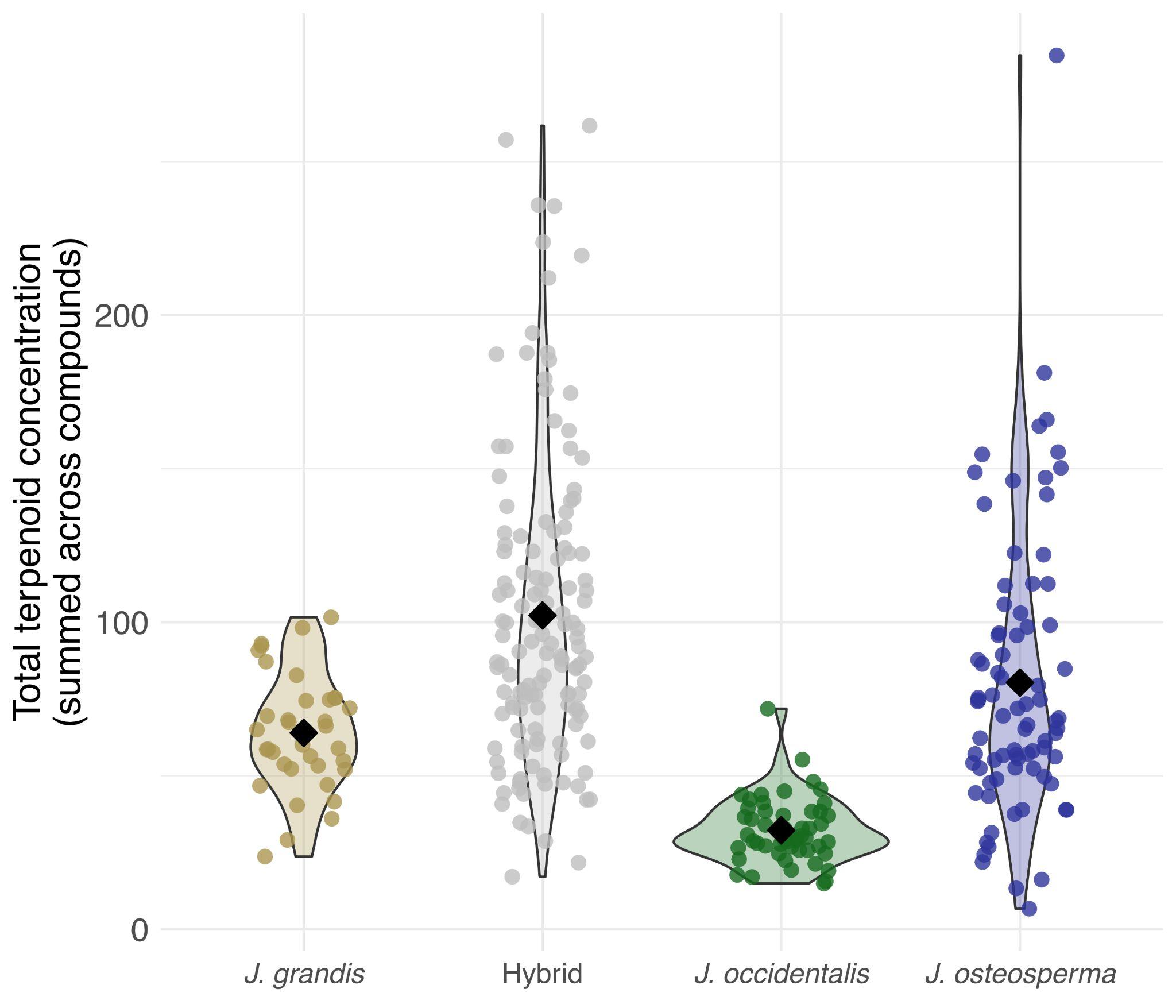

**Figure 5.** Violin plots show the distribution of alpha diversity (effective Shannon diversity) based on terpenoid concentrations within each ancestry group. Alpha diversity was calculated for each sample using the Shannon index. All pairwise comparisons between group means were significant except for *J. osteosperma* and *J. grandis*. Violins are ordered from largest group mean (left) to smallest group mean (right).

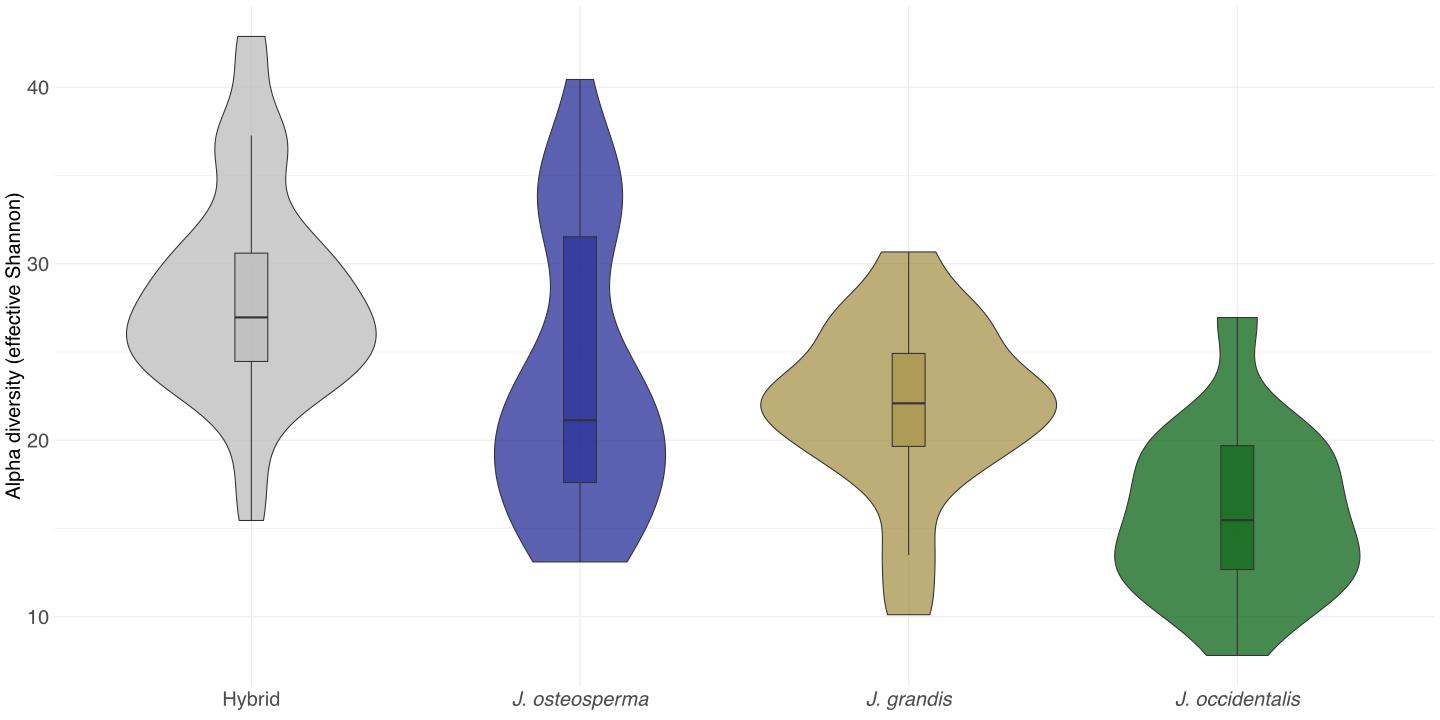

**Figure 6.** Density distributions of the three components of terpenoid beta diversity (β_SOR_, β_SNE_, and β_SIM_) calculated separately for each parental species and the hybrids. β_SOR_ (solid lines) represents overall beta diversity as Sørensen dissimilarity. This is divided into two components: β_SNE_ (dotted-dashed lines), the nestedness component, which reflects terpenoid loss and gain, and β_SIM_ (dashed lines), the turnover component, which captures terpenoid compound replacement among individuals.

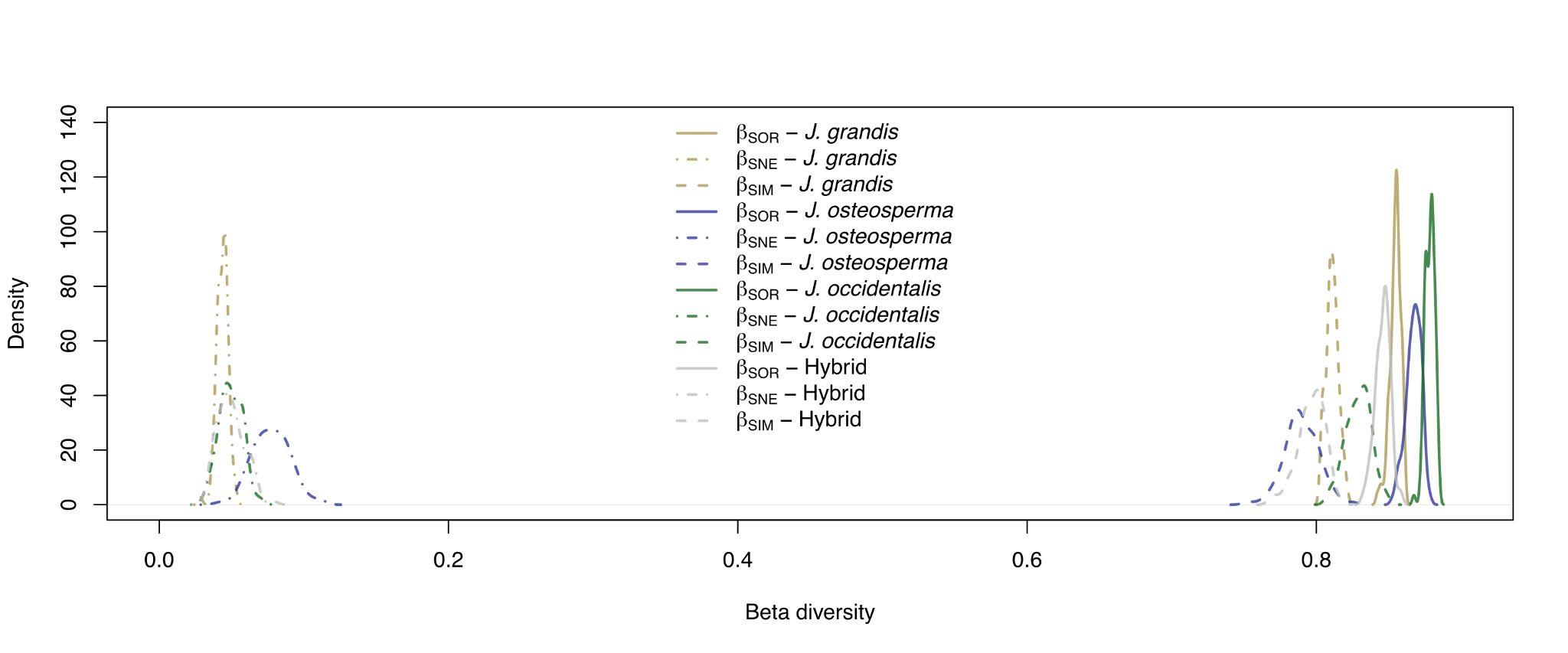

**Figure 7.** Principal Coordinates Analysis (PCoA) plots of terpene profiles, with panels (A–B) displaying Manhattan distances based on terpene concentrations and panels (C–D) showing binary Jaccard distances for terpene presence/absence. Each point represents an individual, with different colors and shapes denoting distinct ancestry groups. Ellipses represent 95% confidence intervals for each group, and larger open points indicate the spatial medians of these groups.

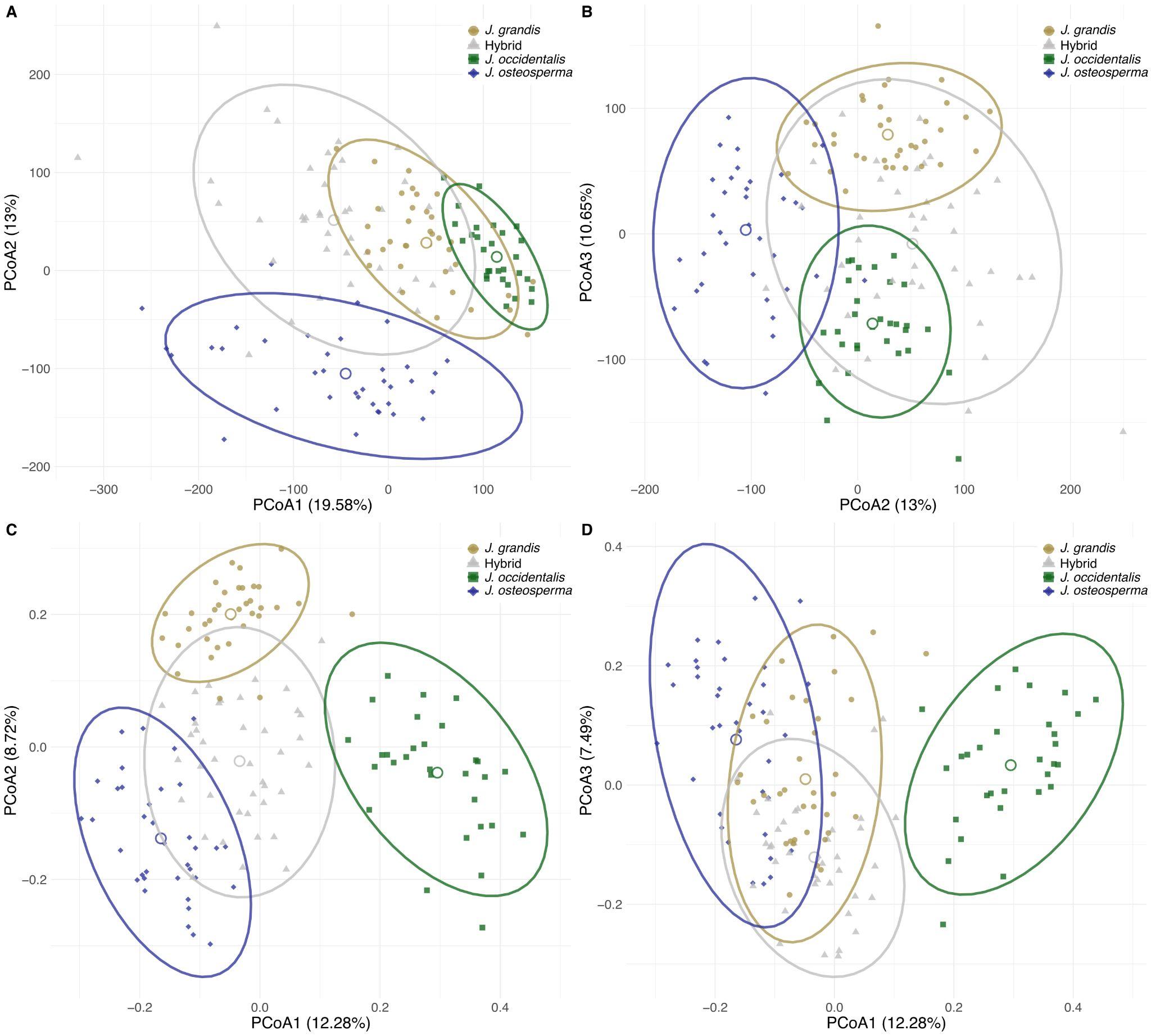
